# Differential expression of five prosomatostatin genes in the central nervous system of the catshark *Scyliorhinus canicula*

**DOI:** 10.1101/823187

**Authors:** Daniel Sobrido-Cameán, Herve Tostivint, Sylvie Mazan, María Celina Rodicio, Isabel Rodríguez-Moldes, Eva Candal, Ramón Anadón, Antón Barreiro-Iglesias

## Abstract

Five prosomatostatin genes (*PSST1*, *PSST2*, *PSST3*, *PSST5* and *PSST6*) have been recently identified in elasmobranchs (Tostivint, Gaillard, Mazan, & Pézeron, 2019). In order to gain insight into the contribution of each somatostatin to specific nervous systems circuits and behaviors in this important jawed vertebrate group, we studied the distribution of neurons expressing *PSST* mRNAs in the catshark *Scyliorhinus canicula* using *in situ* hybridization with specific probes for the five *PSSTs* transcripts. Additionally, we combined *in situ* hybridization with tyrosine hydroxylase (TH) immunochemistry for better localization of some *PSSTs*-positive populations. The five *PSST* genes showed expression in the brain, although with important differences in distribution. *PSST1* and *PSST6* were widely expressed in different brain regions. Instead, *PSST2* and *PSST3* were expressed only in the ventral hypothalamus and in some hindbrain lateral reticular neurons, whereas *PSST5* was only expressed in the region of the entopeduncular nucleus. *PSST1* and *PSST6* were expressed by numerous pallial neurons, although in different populations judging from the colocalization of tyrosine hydroxylase (TH) immunoreactivity and *PSST6* expression in pallial neurons and the absence of colocalization between TH and *PSST1* expression. Differential expression of *PSST1* and *PSST6* was also observed in the subpallium, hypothalamus, diencephalon, optic tectum, midbrain tegmentum and rhombencephalon. Expression of *PSST1* was observed in numerous cerebrospinal fluid-contacting (CSF-c) neurons of the paraventricular organ of the hypothalamus and the central canal of the spinal cord. These wide differences in expression of *PSST* genes together with the numerous brain nuclei expressing *PSSTs*, indicate that catshark somatostatinergic neurons are implicated differentially in a number of neural circuits.

## INTRODUCTION

Somatostatin (SST; also known as somatotropin release inhibiting factor; Brazeau et al., 1974) is a small peptide of 14 amino acids expressed in many brain neuronal populations, as shown in immunohistochemical studies in rats (Vincent et al., 1985). Several isoforms of SST of either 14 amino acids (SST-14) or 28 amino acids (SST-28) have been purified and sequenced from different vertebrates and used for physiological studies showing that SSTs play a key role in the regulation of growth, development and metabolism (see Günther et al., 2018). More recently, different preprosomatostatin genes have been cloned in chondrichthyans (cartilaginous fishes), bony fishes and land vertebrates (Tostivint et al., 1996, 2016; Tostivint, Quan, Bougerol, Kenigfest, & Lihrmann, 2013; Tostivint, Gaillard, Mazan, & Pézeron, 2019). *In situ* hybridization has been employed to study their expression in the brain of some jawed vertebrates (Trabucchi et al., 1999, 2002, 2003, Tostivint et al., 1996).

Elasmobranchs (sharks, rays and skates) are cartilaginous fishes with articulated jaws that diverged over 400 million years ago from the Holocephali (chimaeras), the chondrichthyan group with non-articulated jaws. Because of their phylogenetic position as sister group of Osteichthyes (which contain all established vertebrate model organisms), cartilaginous fishes are a key group for understanding the changes that occurred at the agnathan-gnathostome transition. In sharks, the distribution of SST-like-immunoreactive (-ir) structures has been described in the brain of the leopard shark *Triakis scyllia* (Nozaki, Tsukahara, & Kobayashi, 1984) and the gummy shark *Mustelus manazo* (Chiba, Honma, Ito, & Homma, 1989). Other studies in elasmobranches have also reported the presence of SST-like-ir fibers in the spinal cord (Cameron Plenderleith, & Snow, 1990; Anadón, Molist, Pombal, Rodríguez-Moldes, & Rodicio, 1995) and cerebellum (Alvarez-Otero, Perez, Rodriguez, & Anadón, 1996), and cells and fibers in the hypothalamus (Meurling and Rodríguez, 1990; Molist, Rodriguez-Moldes, & Anadon, 1992). Moreover, in *Squalus acanthias*, SST-like immunoreactivity occurred in peripheral nerve fibers and endocrine cells of the gut (Holmgren and Nilsson, 1983; El-Salhy, 1984). Most physiological studies on SST in elasmobranchs were centered on its action in regulation of salt secretion by the rectal gland (Stoff, Rosa, Hallac, Silva, & Epstein, 1979). More recently, studies on the somatostatinergic system in elasmobranchs have been mainly directed to the identification and sequencing of SST precursor (*PSST*) cDNAs in different species, including the catshark (also called lesser-spotted dogfish) *Scyliorhinus canicula* (Quan, Kenigfest, Mazan, & Tostivint, 2013; Tostivint, Gaillard, Mazan, & Pézeron, 2019). This species is considered as a model of choice in evolutionary developmental biology (Coolen et al., 2009), and has also been the subject of numerous neurochemical and neuroanatomical studies (reviewed in Rodríguez-Moldes, Santos-Durán, Pose-Méndez, Quintana-Urzainqui & Candal, 2017). The three cDNAs previously identified in *S. canicula* were named *ScPSSa*, *ScPSSb* and *ScPSSc* (Quan, Kenigfest, Mazan, & Tostivint, 2013). More recently, Tostivint, Gaillard, Mazan, & Pézeron (2019) identified 2 new *PSST* cDNAs in this species and clarified the five *PSST* phylogeny showing the presence of *PSST1* (previously known in catshark as *ScPSSTa*), *PSST2*, *PSST3*, *PSST5* (previously known in catshark as *ScPSSb*) and *PSST6* (previously known in catshark as *ScPSSTc*) in cartilaginous fishes. The putative mature SST1 and SST5 peptides share the same amino-acid sequence (AGCKNFFWKTFTSC), which is that of the mammalian SST-14. The putative amino-acid sequence of the SST6 mature peptide (A<u>P</u>CKNFFWKTFTSC) differs in position 2 from the mammalian SST (Quan, Kenigfest, Mazan, & Tostivint, 2013; Tostivint, Gaillard, Mazan, & Pézeron, 2019); whereas, the putative amino-acid sequences of the catshark SST2 (<u>TP</u>CK<u>L</u>FFWKTF<u>SH</u>C) and SST3 (<u>N</u>CKNFFWKT<u>Y</u>T<u>L</u>C) mature peptides show more differences in their sequence.

The contribution of specific SSTs to central nervous system (CNS) circuits and behaviors of sharks is completely unknown. To gain insight into the expression of the somatostatinergic system of cartilaginous fishes, we studied the transcript distribution of the PSST paralogs in the CNS of juvenile catshark by means of *in situ* hybridization. Our results reveal that different *PSST* transcripts are expressed differentially in various neuronal populations in this species. These results indicate that the five *PSSTs* are involved in different neuronal circuits and suggest that they have different roles and effects in the CNS. Our study provides a neuroanatomical basis for future functional work on the roles of different SSTs in the CNS of sharks.

## MATERIAL AND METHODS

### Animals

Juveniles (n = 7; between 12 and 19.5 cm in total length) of the catshark *S. canicula* were used for cDNA cloning and *in situ* hybridization experiments. Animals were kindly provided by the Aquarium of *O Grove* (Pontevedra, Spain). Before experimental procedures, fishes were maintained in fresh seawater tanks in standard conditions of temperature (16-18°C), pH (7.5-8.5), salinity (35g/L) and 12:12 h day/night cycle. Adequate measures were taken to minimize animal pain or discomfort. All animal experiments were approved by the Bioethics Committee of the University of Santiago de Compostela and the *Xunta de Galicia* and conformed to the guidelines of the European Communities Council Directive of 22 September 2010 (2010/63/UE) and the Spanish Royal Decree 1386/2018 for the care and handling of animals in research.

### Cloning of the S. canicula SST2 cDNA

A juvenile of the catshark *S. canicula* was anesthetized by immersion in 0.1% ethyl 3-aminobenzoate methanesulfonate salt (MS-222; Sigma, St. Louis, MO, USA) and the brain and spinal cord were dissected out under sterile conditions. Total RNA was isolated from these tissues using TriPure (Roche, Mannhein, Germany). The first-strand cDNA synthesis reaction from total RNA was catalyzed with Superscript III reverse transcriptase (Invitrogen, Waltham, MA, USA) using random primers (hexamers; Invitrogen). For polymerase chain reaction (PCR) cloning, specific oligonucleotide primers (sense 5’-TGGCTGGCTTGTTGGAGACT-3’ and antisense 5’-TGTGGGAAGAGAGGGGGCTA-3’) were designed based on the *catshark PSST2* sequence (Tostivint, Gaillard, Mazan, & Pézeron, 2019). The amplified fragments were cloned into pGEM-T vectors (Promega, Madison, WI, USA) and used for riboprobe synthesis (see below).

### In situ hybridization

For the generation of the catshark *PSST1, PSST3, PSST5* and *PSST6* riboprobes we used a *S. canicula* embryonic cDNA library available in Dr Mazan’s laboratory (see Quan, Kenigfest, Mazan, & Tostivint, 2013; Tostivint, Gaillard, Mazan, & Pézeron, 2019). For the generation of the catshark *PSST2* riboprobe we used the *PSST2* clone that we obtained (see above). DIG-labeled riboprobes for shark *PSSTs* were synthesized also by *in vitro* transcription.

Juvenile catsharks were deeply anesthetized with 0.1% ethyl 3-aminobenzoate methanesulfonate salt (MS-222; Sigma, St. Louis, MO, USA) in seawater before experimental procedures. *In situ* hybridization experiments were performed as previously described for riboprobes against the sea lamprey serotonin 1a receptor mRNA (Cornide-Petronio, Anadón, Barreiro-Iglesias, & Rodicio, 2013). Briefly, catsharks (n = 6) were perfused intracardially with elasmobranch Ringer’s solution (see Ferreiro-Galve, Rodríguez-Moldes, & Candal, 2012) followed by 4% paraformaldehyde in elasmobranch’s phosphate buffer (0.1 M phosphate buffer containing 1.75% urea, pH 7.4). Catshark brains and rostral spinal cords were dissected out and post-fixed in the same fixative for 48 h at 4 °C. Then, they were cryoprotected with sucrose 30% and sectioned on a cryostat in the transverse plane (14 µm sections). Five parallel series of sections were obtained from each brain/spinal cord block. The sections of each series were incubated with each of the 5 *PSST* DIG-labelled probes, respectively, at 70°C overnight in hybridization mix and treated with RNAse A (Invitrogen, Waltham, MA, USA) in the post-hybridization washes. Then, the sections were incubated with sheep anti-DIG antibodies conjugated to alkaline phosphatase (1:2000; Roche, Mannheim, Germany) overnight. Staining was conducted in BM Purple (Roche) at 37°C until the signal was clearly visible. Finally, the sections were mounted in Mowiol (Calbiochem; Temecula, CA, USA).

### Double *in situ* hybridization and immunohistochemistry

In some samples, *in situ* hybridization for *PSST1* and *PSST6* was followed by immunohistochemistry against tyrosine hydroxylase (TH). After the *in situ* hybridization signal was clearly visible, sections were rinsed twice in 0.05 M Tris-buffered saline (TBS; pH 7.4) and treated with 10% H_2_O_2_ in TBS for 30 minutes. For heat-induced epitope retrieval, sections were treated with 0.01 M citrate buffer (pH 6.0) for 30 min at 90°C and allowed to cool for 20–30 min at room temperature (RT) in the same buffer. Then, the sections were incubated in primary antibody solutions overnight at RT. The primary antibody was a mouse monoclonal anti-TH antibody (a mouse monoclonal antibody raised against TH purified from PC12 cells; Millipore, CA; Cat#MAB318; RRID: AB_2201528; dilution 1:1,000; Table 1). After rinsing in TBS, the sections were incubated with the solution containing the secondary antibody (goat anti-mouse IgG serum HRP conjugated (Dako, Glostrup, Denmark; Cat# G-21040, RRID: AB_2536527, dilution 1:200; Table 1) for 1 hour at RT. All antibodies were diluted in TBS containing 15% normal serum of goat and 0.2% Triton X-100 (Sigma). The immunoreaction was developed with 0.25 mg/ml diaminobenzidine tetrahydrochloride (DAB; Sigma) containing 0.00075% H_2_O_2_. Finally, the sections were mounted in Mowiol.

**Table 1.**
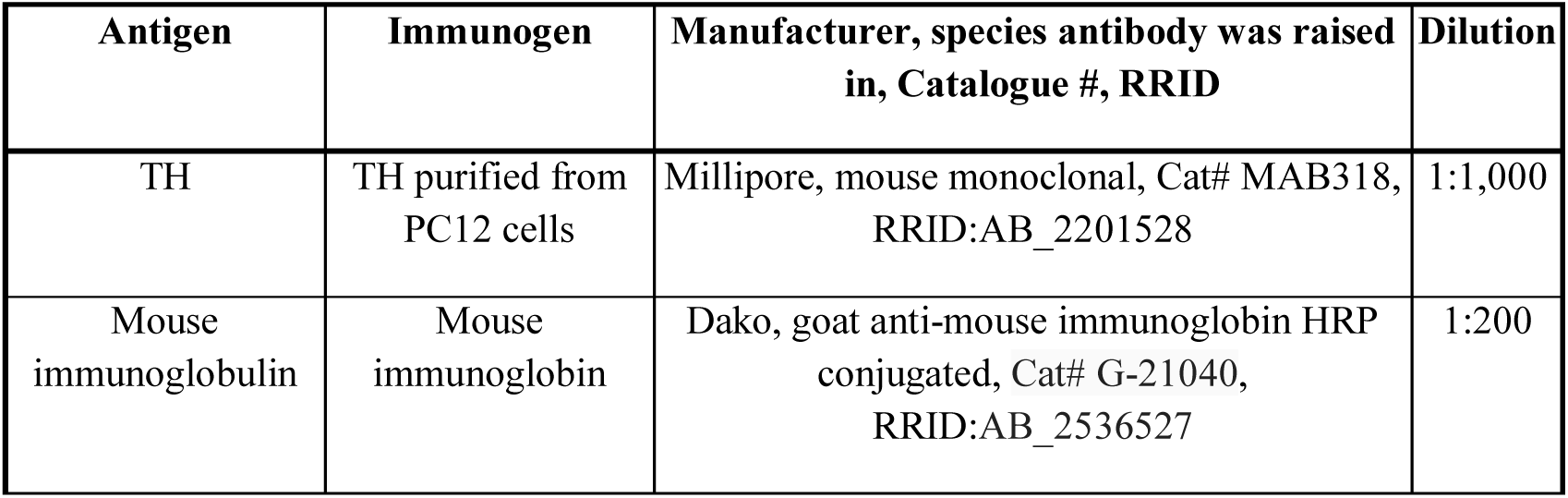
Antibodies used in this study.

| Antigen | Immunogen | Manufacturer, species antibody was raised in, Catalogue #, RRID | Dilution |
| --- | --- | --- | --- |
| TH | TH purified from PC12 cells | Millipore, mouse monoclonal, Cat# MAB318, RRID:AB_2201528 | 1:1,000 |
| Mouse immunoglobulin | Mouse immunoglobulin | Dako, goat anti-mouse immunoglobulin HRP conjugated, Cat# G-21040, RRID:AB_2536527 | 1:200 |

### Primary antibody characterization

The mouse monoclonal TH antiserum was raised against denatured TH from rat pheochromocytoma. According to the technical information supplied by the manufacturer, it recognizes an epitope on the outside of the regulatory N-terminus and in Western blots it recognizes a protein of approximately 59-61 kDa. In addition, the antiserum specificity was characterized by Western blot in brain extracts of the catshark, in which it stained a single protein band of about 56-60 kDa (Carrera et al., 2012). The antibody displays wide species cross-reactivity and has been used to demonstrate the catecholaminergic systems in a number of species, including the catshark (reviewed in Carrera, Anadón, & Rodríguez-Moldes, 2012). The antibody does not recognize TH2 in teleosts, revealing only the TH1-immunoreactive catecholaminergic neurons (Filippi, Mahler, Schweitzer, & Driever, 2010; Yamamoto, Ruuskanen, Wullimann, & Vernier, 2010).

### Imaging

A photomicroscope (Provis AX-70; Olympus, Tokyo, Japan) equipped with a color digital camera (Olympus DP70, Tokyo, Japan) was used to acquire images of brain and spinal cord sections. Contrast and brightness of photomicrographs were minimally adjusted with Adobe Photoshop CS4 (Adobe Systems, San Jose, CA, USA). Figure plate composition and lettering were generated using Adobe Photoshop and schematic drawings were made using CorelDRAW 12 (Corel, Ottawa, Canada).

### Nomenclature for brain structures

For description of brain nuclei in the juvenile catshark brain we followed in general the nomenclature of Smeets, Nieuwenhuys, & Roberts (1983) adapted to new ideas on the segmental organization of the catshark brain, including changes in the limits between the major brain segments based on developmental and genoarchitectonic studies (Anadón et al., 2000; Carrera et al., 2005; Carrera, Molist, Anadón, & Rodríguez-Moldes, 2008a; Carrera, Ferreiro-Galve, Sueiro, Anadón, & Rodríguez-Moldes, 2008b; Carrera, Anadón, & Rodríguez-Moldes, 2012; Rodríguez-Moldes et al., 2017; see Table 2).

**Table 2.**
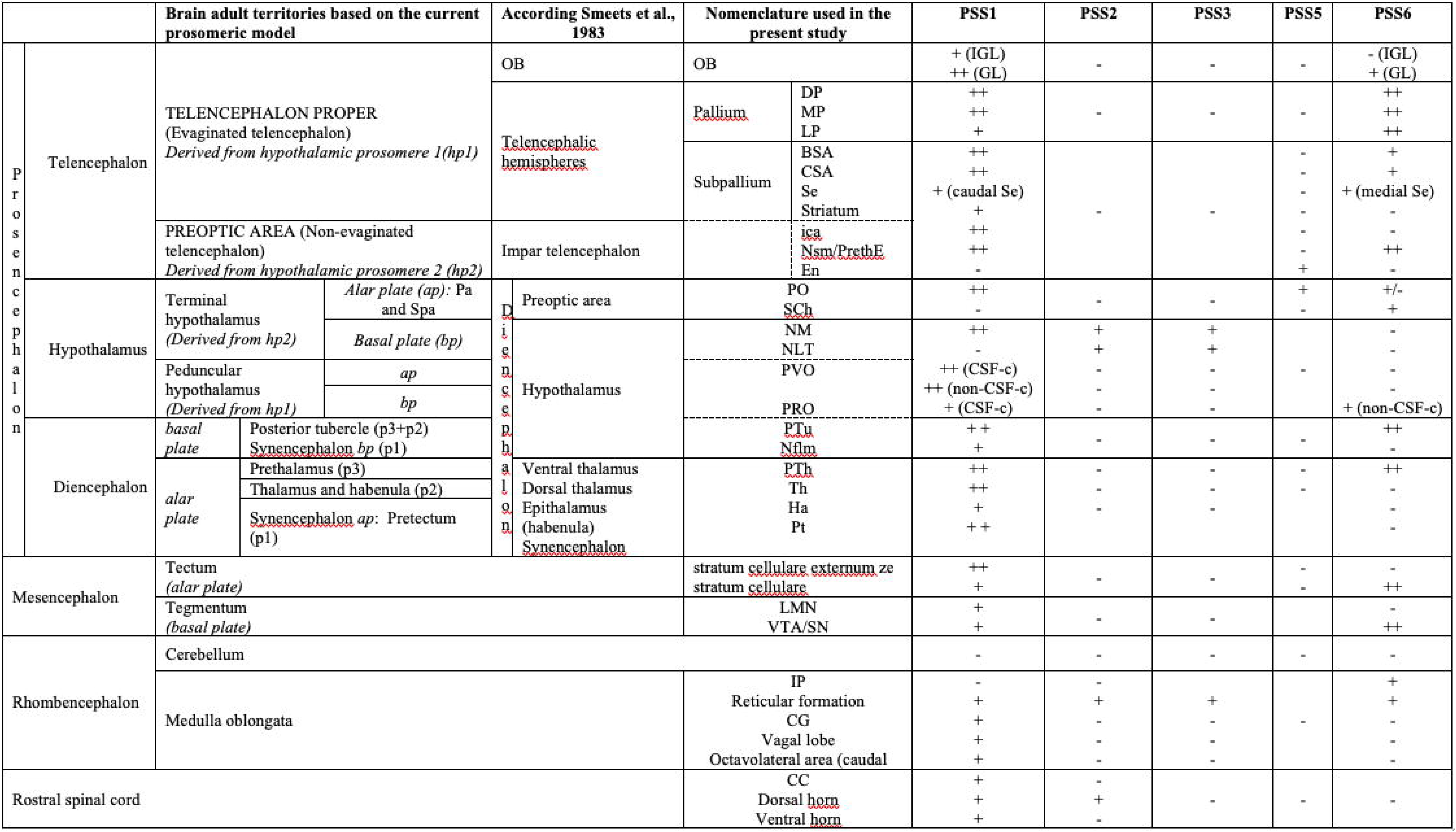
Expression PSST transcripts in cells of the major brain components and rostral spinal cord of juvenile catshark. The correspondence of juvenile territories according the current prosomeric model and the nomenclature of Smeets, Nieuwenhuys, & Roberts (1983) is depicted. For abbreviations, see list. –, no expression; +, some expression; ++, abundant expression.

## RESULTS

*In situ* hybridization of sections of the brain and rostral spinal cord of juvenile catshark showed differential expression of *PSST* genes. Two of these genes (*PSST1* and *PSST6*) showed wide expression in the brain (Fig. 1-6, Table 2), whereas expression of *PSST2*, *PSST3* and *PSST5* was limited to a few brain structures (Fig. 7 and 8, Table 2). In the rostral spinal cord, only *PSS1* and *PSS2* were expressed. For the description of the *PSST*-expressing neuronal populations we follow a rostro-caudal sequence. As far as possible, the nomenclature for brain regions is updated taken into account the new studies on regional brain organization of the catshark (e.g. Rodríguez-Moldes, Santos-Durán, Pose-Méndez, Quintana-Urzainqui, & Candal, 2017; Table 2). In currently available adult catshark brain maps, however, there are extensive “reticular” areas that lack well-characterized nuclei but exhibit various somatostatinergic populations. These “reticular” populations will be referred here according its segmental position, its presumed alar or basal location, and its location relative to other topological references. In order to assess the location of some somatostatinergic populations, we also used double staining with *in situ* hybridization for *PSST1* or *PSST6* and immunohistochemistry for TH. TH is a general marker of catecholaminergic neuronal populations whose distribution in the brain of elasmobranchs including catshark has been well characterized (Northcutt, Reiner, & Karten, 1988; Stuesse, Cruce, & Northcutt, 1994; Carrera et al., 2005; Carrera, Anadón, & Rodríguez-Moldes, 2012).

**Figure 1.**
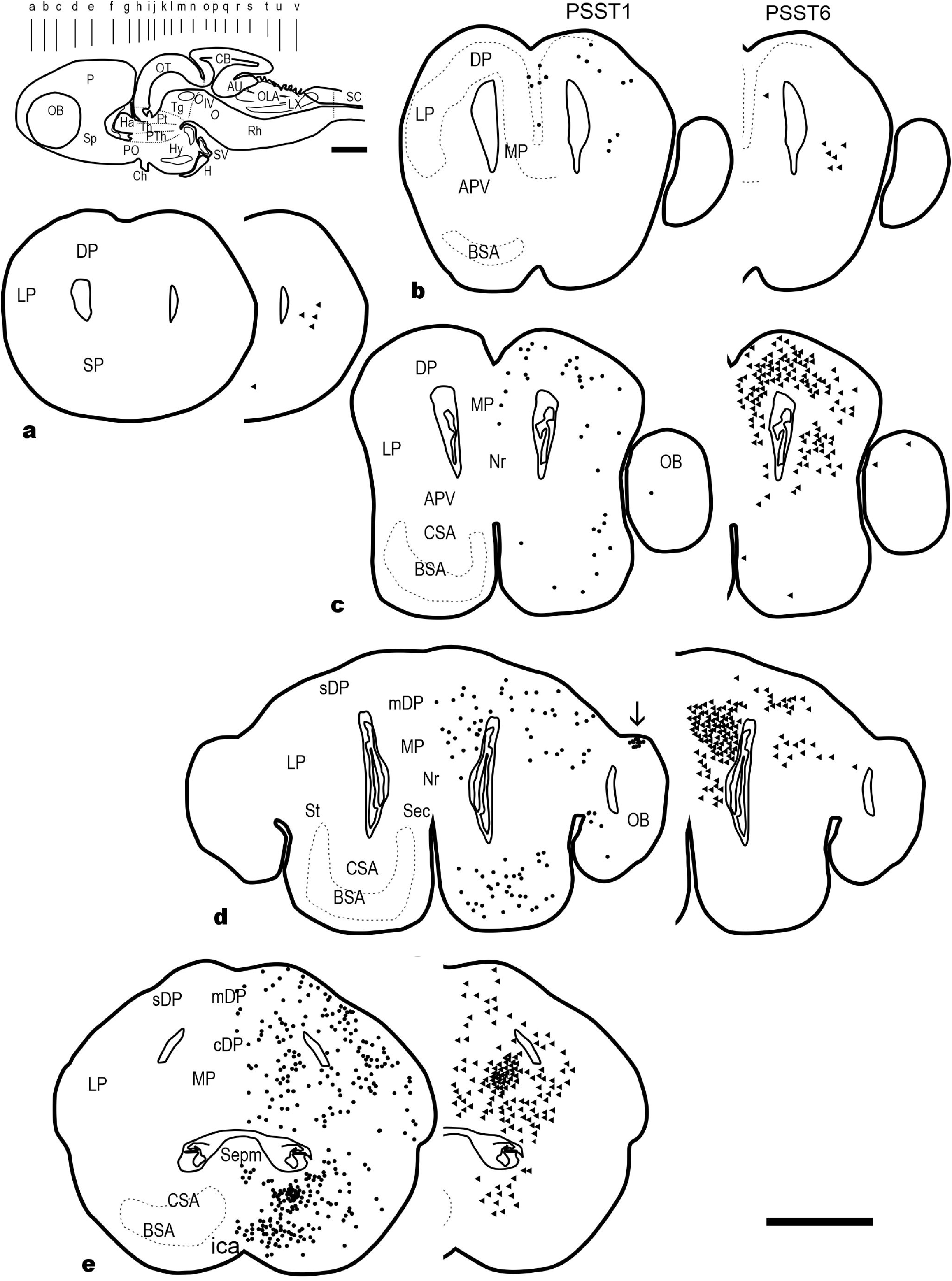
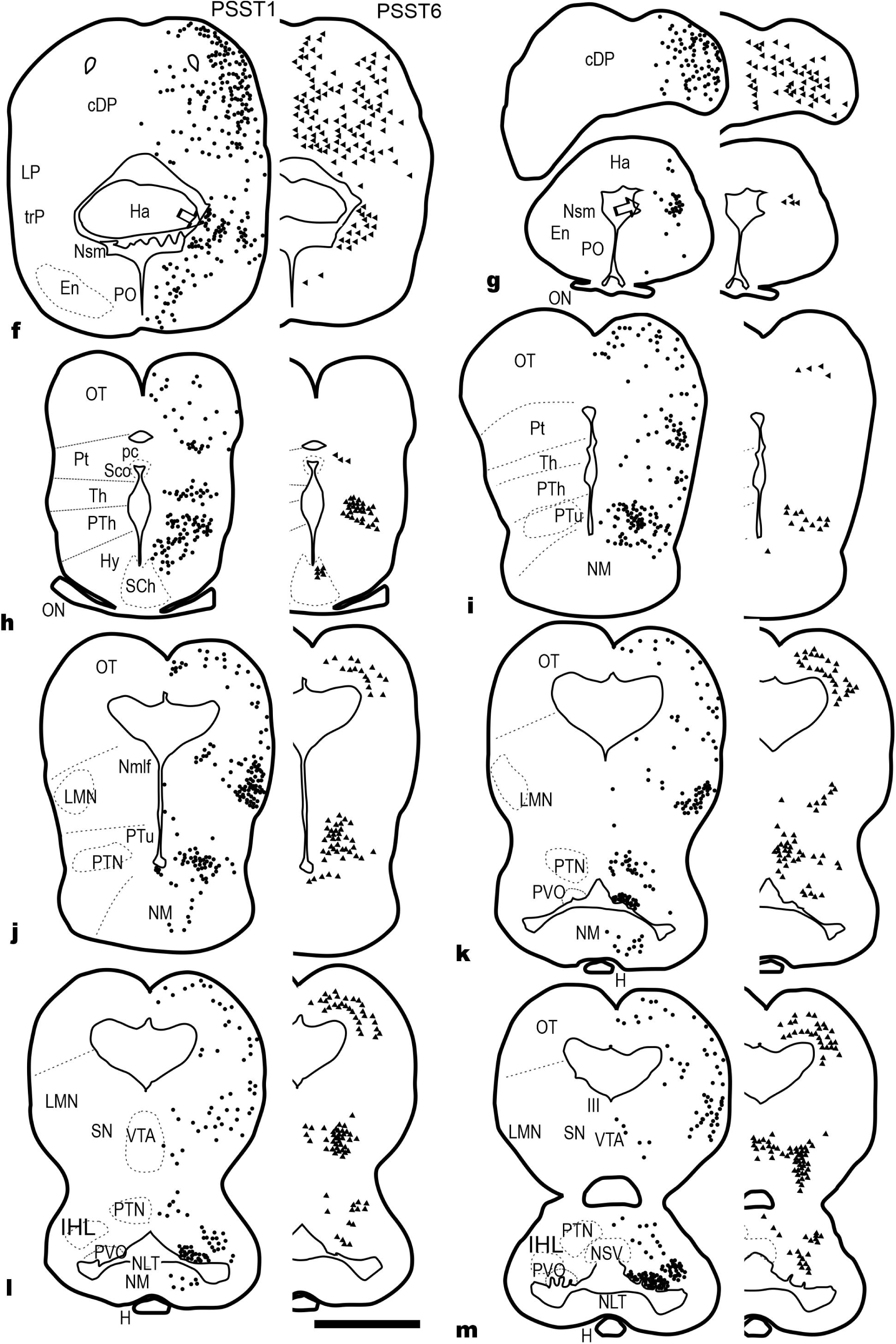
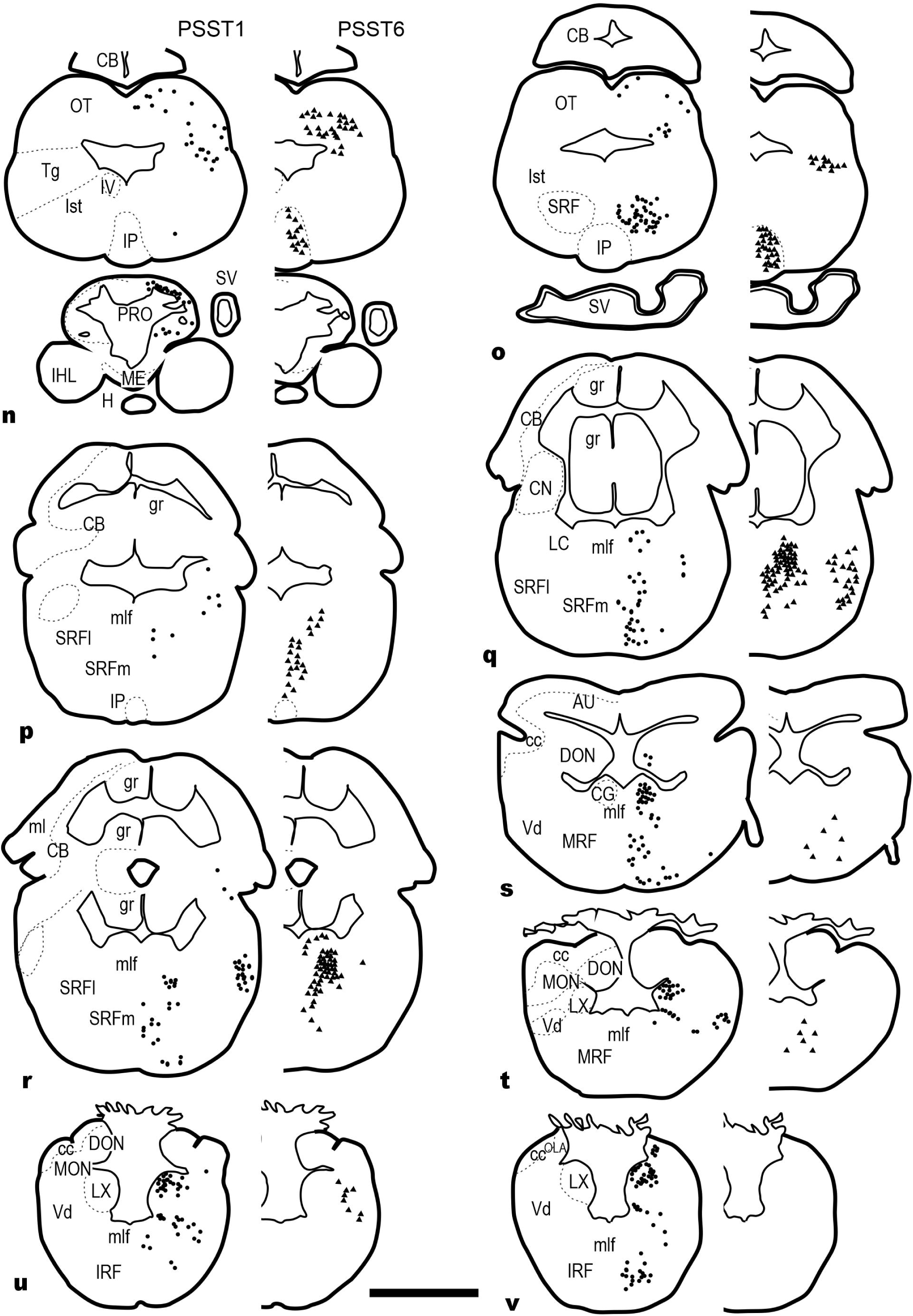
Schematic drawings of parallel transverse sections of a juvenile brain *in situ* hybridized for *PSST1* or *PSST6* showing the plotted distribution of cells expressing *PSST1* mRNA (dots in the left figurines) and *PSST6* mRNA (black triangles in the right figurines). Each symbol corresponds to a positive cell in a 16µm-thick section. Anatomical structures are indicated in the left side of the left figurines. For abbreviations, see the list. The arrow in **d** points to a compact group of *PSST1*+ cells in the olfactory bulb. The level of sections is indicated at the upper left corner in a schematic drawing of a juvenile brain. Scale bars: 1 mm.

### *PSST1* expression

#### Telencephalon

The telencephalon is considered to be an evaginated alar derivative of the secondary prosencephalon, which also includes the preoptic region and the hypothalamus (Rodríguez-Moldes, Santos-Durán, Pose-Méndez, Quintana-Urzainqui, & Candal, 2017). The large catshark telencephalon has paired hemispheres that are widely confluent in the midline producing a massive appearance in juveniles and adults (Smeets et al., 1983; Rodríguez-Moldes, Santos-Durán, Pose-Méndez, Quintana-Urzainqui, & Candal, 2017). The lateral ventricles are reduced at most parts to curved flattened canals that join caudally to the impar telencephalic ventricle. Big olfactory bulbs protrude laterally from the hemispheres. Caudally the impar telencephalon joins it with the preoptic region. The hemispheres arise from two main histogenetic subdivisions, the pallium and the subpallium. The catshark developing pallium may be subdivided in at least four different histogenetic territories: medial, dorsal, lateral and ventral by using early developmental gene markers (Rodríguez-Moldes, Santos-Durán, Pose-Méndez, Quintana-Urzainqui, & Candal, 2017). In the subpallium, the putative striatal and pallidal homologues have been defined using patterns of early developmental gene expression in embryos (Quintana-Urzainqui et al., 2012, Rodríguez-Moldes, Santos-Durán, Pose-Méndez, Quintana-Urzainqui, & Candal, 2017). Based on such studies, the preoptic area was recognized as a subpallial subdomain. Equivalences have also been proposed between these embryonic primordia and adult subpallial territories, although some uncertainties exist. Since gene expression-based correspondence among the pallial and subpallial subregions of adults with those of other vertebrates was not unequivocally established, for the cell masses of juvenile catshark telencephalon we will use classical subdivisions (Smeets, Nieuwenhuys, & Roberts, 1983; Manso and Anadón, 1993) based on the topography with respect the ventricles. Equivalences with developmental data will be also mentioned when appropriate (see Table 2).

The most anterior region of the catshark pallium, which borders rostrally the lateral ventricles, lacks *PSST1* positive (+) cells. Caudally, small *PSST1*+ cells appear progressively mainly in intermediate regions of the pallium (dorsal, medial and lateral to the ventricle). In intermediate and caudal regions these cells appear distributed from periventricular to superficial layers, and *PSST1*+ cells are particularly abundant in the most caudolateral region of the pallium (dorsal pallium of Smeets et al., 1983) (Figs. 1a-g, 2a-d). The pallium shows a fairly abundant population of small TH-immunoreactive (TH-ir) cells that are *PSST1* negative (Figs. 2b-d, 3f).

In the rostral subpallium, at levels showing the thick cell band of the conspicuous basal superficial area (BSA), small and faint *PSST1+* cells are located scattered among negative cells as well as in the adjacent outer and inner cell and neuropil areas. These BSA cells are scarce rostrally but their number increases notably toward caudal regions (Figs. 1c-e, 2a-b). Caudally, numerous intense *PSST1*+ cells are distributed in the wide basal central area of cells and neuropil between the BSA and the ventricles (Fig. 1e, 2b). These larger cells are conspicuous in caudal levels near or within the BSA. The transition of the BSA to the interstitial nucleus of the anterior commissure is marked by the appearance of a population of small TH-ir cells in this nucleus and in the caudal septum, whereas the BSA is free of these cells along its rostrocaudal extension. Scattered *PSST1*+ cells were observed in the caudal septum of Smeets, Nieuwenhuys, & Roberts (1983); whereas, abundant *PSST1+* cells were observed in the region referred as posterior medial septum by these authors (Figs. 1e, 2b). Ventromedial to the caudal olfactory bulb and lateral to the BSA there is a region of scattered cells named as striatum by Smeets, Nieuwenhuys, & Roberts (1983). This ill-defined striatal region shows scarce *PSST1*+ cells compared with the BSA and adjacent areas.

**Figure 2.**
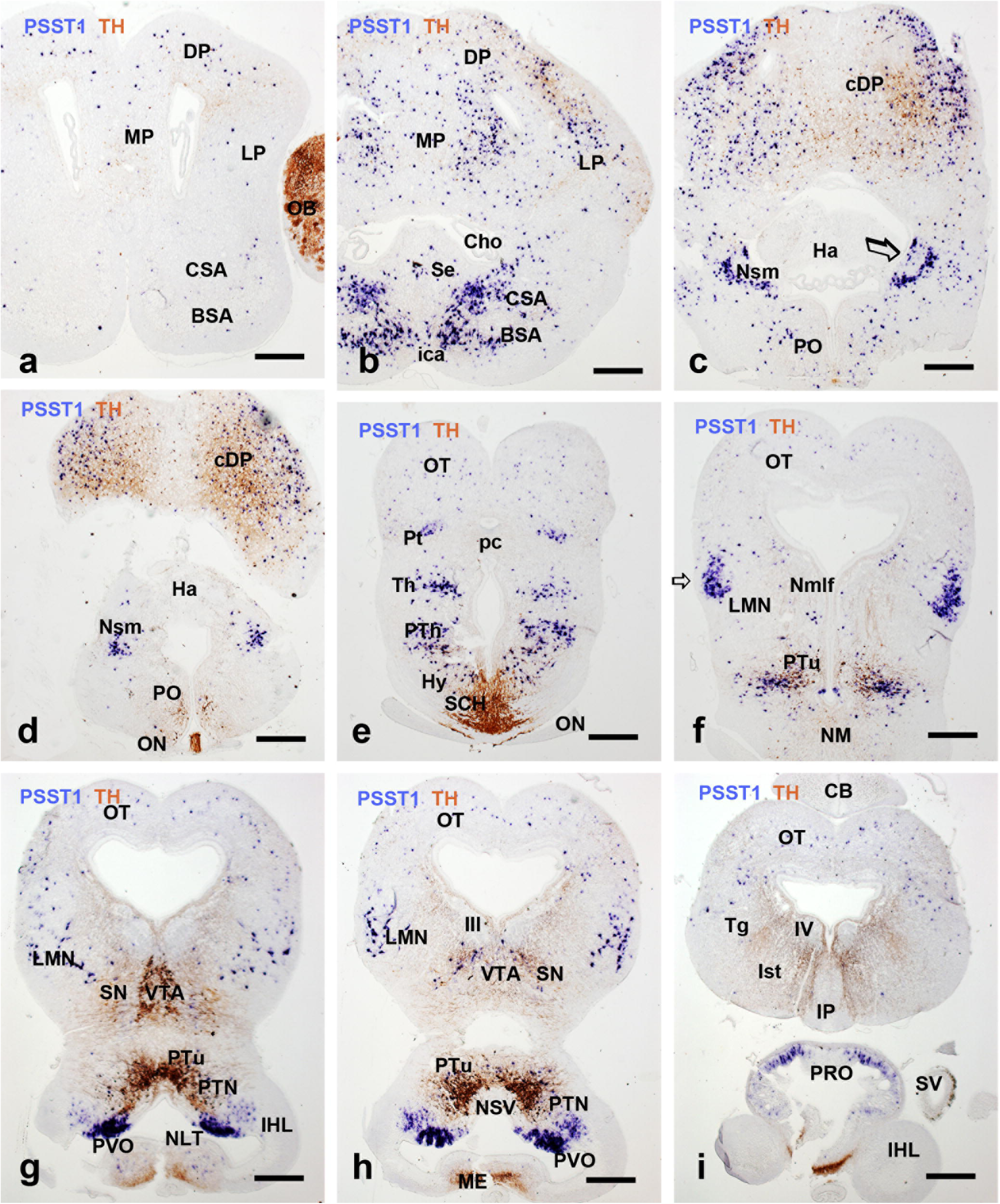
Low magnification photomicrographs of double stained transverse brain sections of a juvenile catshark showing in blue *PSST1* positivity and in brown TH immunoreactivity. These sections correspond to the brain shown schematically at the left side in Figure 1. For abbreviations, see the list. Scale bars, 250 µm.

At the transition with the telencephalic peduncle, a dense group of small *PSST1*+ cells is observed along the region of the stria medullaris and in a small protuberance close to the insertion of the anterior medullary velum (Figs. 1f-g, 2c-d, arrowed). This group ends close to the habenula and may be considered a part of the prethalamic eminence (nucleus of the stria medullaris). The interstitial nucleus of the anterior commissure shows numerous small *PSST1+* cells (Figs. 1e, 2b). This population ends in the rostral preoptic region.

The olfactory bulbs of the juvenile catshark protrude laterally from the lateral pallial region (Fig. 1c-d). Using double *PSST1* and TH staining, a large population of small TH-ir cells is observed close and between the negative large mitral cells (Fig. 2a, 3g). Processes of these TH-ir cells help to visualize the large olfactory glomeruli (not shown). In addition to these TH-ir periglomerular cells, scattered TH-ir cells are observed in the inner granular layer and periventricular regions of the olfactory bulbs (see Carrera, Anadón, & Rodríguez-Moldes, 2012). In clear contrast with the prominent distribution of TH immunoreactivity, only a few *PSST1*+ cells are observed per section in the OB, mostly in the inner granular layer and in caudal regions of the OB. However, a small compact group of *PSST1* + neurons was observed in the dorsal outer border of the glomerular layer, which lacks TH staining (Figs. 1d, 3g).

**Figure 3.**
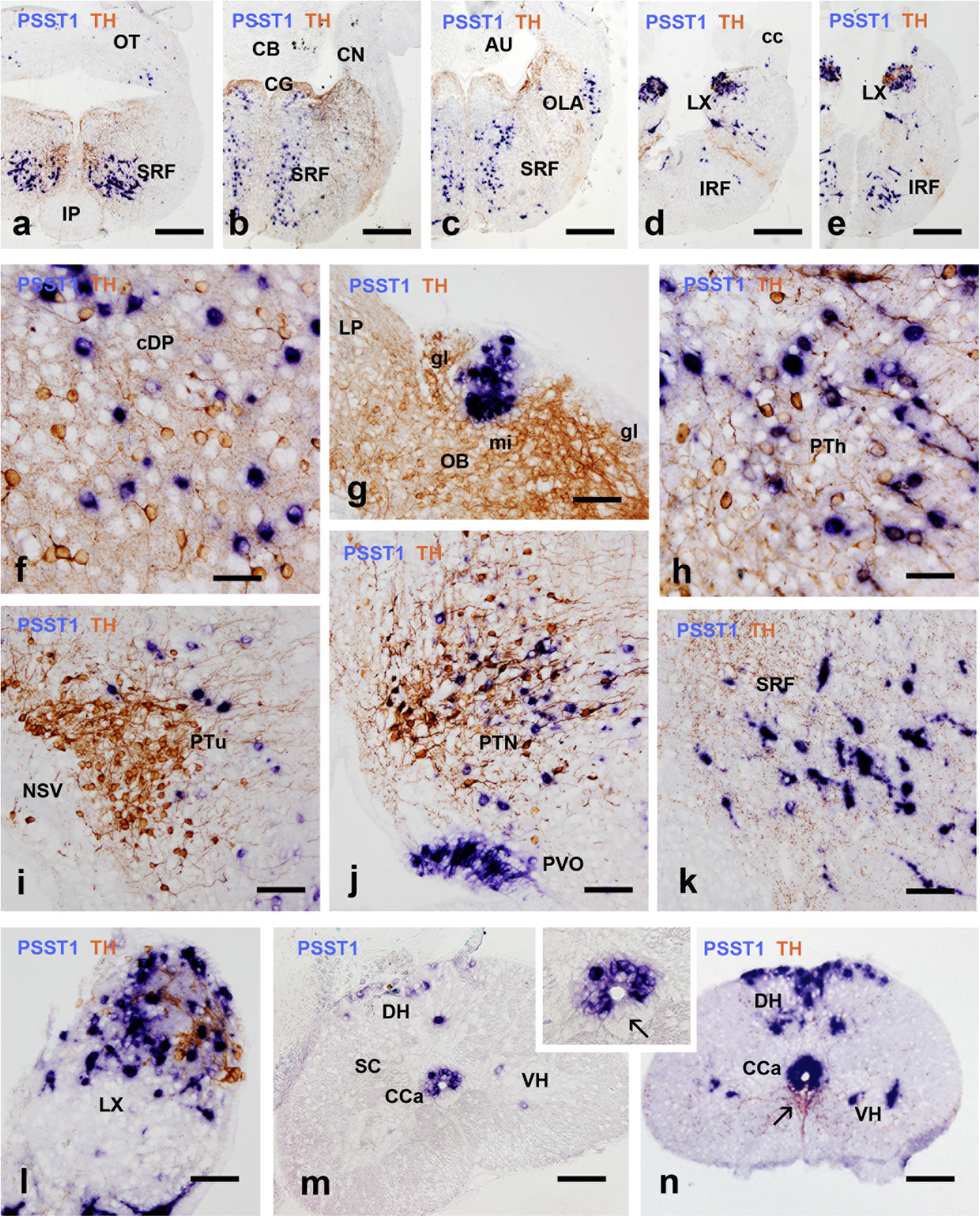
Photomicrographs of double stained transverse brain sections of a juvenile catshark showing *PSST1* positivity in blue and TH immunoreactivity in brown. **a-e**, Low magnification photomicrographs of the hindbrain. The level of these brain sections roughly corresponds to that shown schematically at the left side in Figure 1. **f-l**, Details of sections showing co-distribution of *PSST1*+ and TH-ir structures. **m-n**, sections of the spinal cord showing *PSST1* in situ hybridization alone (**m**) or combined with TH-ir staining (**n**). Inset is a detail of the *PSST1*+ cells around the central canal excepting a ventral sector that only shows TH-ir CSF-c cells (arrows). For abbreviations, see the list. Scale bars, 250 µm (a, b, c, d, e), 100 µm (g, i, j, k, m, n), 50 µm (f, h, l, inset in m).

#### Preoptic region and hypothalamus

Here we consider as the preoptic region the territory of cells and fibers that extends laterally to the preoptic recess and rostral to the optic chiasm. This region joins the subpallium dorsally and the alar hypothalamus ventrally (in a topological view). At the level of the optic chiasm a dense population of TH-ir cells forms part of a prominent suprachiasmatic nucleus (alar hypothalamus) that is partially imbricated with the TH-negative optic nerve fascicles (Fig. 2d-e). A few *PSST1*+ cells are observed in preoptic and suprachiasmatic lateral regions, whereas, more numerous TH-ir neurons are located in the periventricular regions (Fig. 2d-e).

In the hypothalamus, strong *PSST1*+ expression was observed in numerous cells of the hypothalamic circumventricular (vascular) organs. These structures are located in the walls of ventral infundibular recess and extends from levels just rostral to the lateral hypothalamic recesses, invading there a band of small recesses extending laterally in its dorsal wall (hypothalamic paraventricular organ, PVO) and reaching the posterior recess walls, where they form part of the posterior recess organ (PRO) (Figs. 1k-n, 2g-i, 3j). Judging from the bipolar appearance of positive cells and its location inside the ependymal layer (see Fig. 3j), most of them probably correspond to cerebrospinal fluid-contacting (CSF-c) cells. In the posterior recess organ, the distribution of these *PSST1*+ cells is sparser than in the lateral recesses. Accompanying these circumventricular organs, there is a population of faint *PSST1*+ cells, probably non-CSF-contacting, that extend latero-dorsally to the diverticles of the lateral recess (Figs. 1-l-m, 2g-h). In the median hypothalamic nucleus, below the lateral tuberal nucleus of Smeets, Nieuwenhuys, & Roberts (1983), there is a sparse population of faint *PSST1*+ cells (Figs. 1k-l, 2f).

#### Diencephalon

The catshark diencephalon is a region intermediate between the secondary prosencephalon (telencephalon plus hypothalamus) and the mesencephalon. According the evolving prosomeric model (Puelles and Rubenstein, 2003), the embryonic vertebrate diencephalon is formed by three transverse brain segments or prosomeres p1-p3 (from caudal to rostral), whose main alar derivatives are the pretectum, the thalamus (formerly named dorsal thalamus) plus the habenula and the prethalamus (formerly the ventral thalamus), respectively. Main basal derivatives of these prosomeres include the posterior tubercle and the nucleus of the medial longitudinal fascicle.

In the prethalamus-posterior tubercle (p3), a group of *PSST1*+ cells extends in lateral regions (Figs. 1h-m, 2e-f, 3h-j). This group is more compact dorsally than in the posterior tubercle. Except in most rostral (dorsal) prethalamic regions, cells of this *PSST1*+ subpopulation are intermingled with tubercular TH-ir cells, but colocalization of both substances was not observed (Fig. 3h-j). Limits with dorsal hypothalamic *PSST1*+ populations are not sharp, but the diencephalo-hypothalamic external sulcus (also named sulcus thalamohypothalamicus, Smeets, Nieuwenhuys, & Roberts, 1983) can be used as a reference.

In the thalamus, there is an intermediate population of *PSST1*+ cells between the periventricular cell band and the meninges. This population is more condensed rostrally and becomes scattered caudally (Figs. 1h-i, 2e).

In the most dorsal pretectal region, a compact group of *PSST1*+ cells is observed laterally among the fibers of the posterior commissure tract (Figs. 1h, 2e). Caudally this group extends towards the lateral surface below the optic tectum and expands in a bulge of the superficial pretectum (Fig. 2f, arrowed), which shows a large population of small *PSST1*+ cells. Probably this bulge corresponds to the region formerly named as “nucleus geniculatus lateralis” (Smeets, Nieuwenhuys, & Roberts, 1983). In addition to this superficial group, small *PSST1*+ cells are observed scattered between this group and the caudal periventricular region.

#### Mesencephalon (midbrain)

The catshark midbrain is easily identifiable by the presence of a characteristically layered optic tectum dorsally (Fig. 1h-o) and a wide tegmental region ventrally. The midbrain contains a ventricle, showing a hearth-like appearance at middle levels. Whereas the dorsal limits with the pretectum rostrally and the anterior medullary velum caudally are clear in transverse sections, the limits of the midbrain tegmentum with the diencephalon and the isthmus and the ascription of some somatostatinergic populations are more difficult to recognize, especially in lateral regions.

The catshark optic tectum has been subdivided into six layers of neurons and fibers, named from the meningeal to the ependymal layer as stratum medullare externum (optic fiber layer), stratum cellulare externum zona externa, stratum cellulare externum zona interna, stratum medullare internum, stratum cellulare internum and stratum periventriculare (Smeets, Nieuwenhuys, & Roberts, 1983; Manso and Anadón, 1991a). Numerous faintly stained *PSST1*+ small cells are scattered mainly through the stratum cellulare externum zona externa, but sparse *PSST1+* cells were also observed in the zona interna and occasional cells in the stratum cellulare internum (Figs. 1h-o, 2e-i).

In the midbrain tegmentum, the most conspicuous *PSST1*+ population was observed in the lateral mesencephalic nucleus (Figs. 1j-l, 2f-h). It consists of strongly stained cells located laterally near the meninges, and ventrally to the optic tectum. Faint *PSST1*+ cells were also observed intermingled with TH-ir neurons in the ventral tegmental area and the substantia nigra (Figs. 1m, 2g-h), but they do not form a conspicuous group as that of *PSST6*+ cells (see below). Faint *PSST1*+ small cells were also observed scattered at the transition between the optic tectum and the tegmentum.

#### Rhombencephalon (hindbrain)

Just caudal to the border with the midbrain tegmental region, a conspicuous population of strongly *PSST1*+ reticular cells appears in intermedio-lateral regions of the superior reticular formation at the level of the interpeduncular nucleus, which lacks *PSST1*+ cells (Figs. 1o, 3a,k). This conspicuous *PSST1*+ population disappears at intermediate rostrocaudal levels of the interpeduncular nucleus.

More caudally, at the level of the locus coeruleus (which contains TH-ir cells; Carrera, Anadón, & Rodríguez-Moldes, 2012), groups of small *PSST1*+ cells were observed extending in a vertical band near the midline between the central grey and the ventral surface, and a few cells also were present in the central grey (Figs. 1q, 3b-c), probably forming part of the raphe and the superior reticular formation, which also contains serotonergic neurons (Carrera, Molist, Anadón, & Rodríguez-Moldes, 2008a). More caudally, some ventral cells of these groups extend laterally forming a sparse band parallel to the meninges (Fig. 1r-s) appearing as migrating/migrated cells similarly to reticular serotoninergic neurons of the catshark superior reticular formation (Carrera, Molist, Anadón & Rodríguez-Moldes, 2008a). Caudally, the *PSST1+* vertical band diminishes gradually its number of cells. At the level of the trigeminal nerve entrance, a group of strongly *PSST1*+ cells appears over the medial longitudinal fasciculus. Both populations extend caudally until the level of the octaval nerve entrance.

At the level of the facial nerve entrance, *PSST1*+ cells appear around the solitary fascicle, which occupies a small longitudinal protuberance just dorsal to the sulcus limitans that is continuous with the vagal lobe. The number of *PSST1*+ cells is small before the lobe but increases notably in the vagal lobe (Figs. 1t-v, 3d-e, l). In this lobe, the distribution of *PSST1*+ cells, as that of TH-ir cells observed in this region (Carrera. Anadón, & Rodríguez-Moldes, 2012; present results) is dorsal, with approximately the ventral half of the lobe free of these cells (Fig. 3l). Double staining reveals that *PSST1+* and TH-ir cells represent different populations. A conspicuous band of strong *PSST1*+ reticular cells extending ventrolaterally from the sulcus limitans region accompanies most of the extension of the solitary tract-vagal lobe (Fig. 1t-v).

At caudal hindbrain levels, a conspicuous population of large *PSST1*+ reticular neurons is observed in the inferior reticular formation (Figs. 1v, 3e). These cells show bipolar or triangular appearance, with one or more stained dendritic shafts. A few *PSST1*+ cells were observed in the caudal region of the octavolateralis area (OLA, Fig. 1v) and in a lateral group of the rostral OLA (Fig. 3c).

#### Spinal cord

In the rostral spinal cord, a conspicuous population of *PSST1*+ neurons was observed around the central canal except in its ventral side (Fig. 3m). The position and location is compatible with that of CSF-c neurons of the central canal. Double staining reveals that CSF-c TH-ir cells occupy the ventral region of the central canal free of *PSST1*+ neurons (Fig. 3n). In addition to putative CSF-c cells, larger *PSST1*+ cells were observed scattered in the dorsal and ventral horns (Fig. 3m-n).

### *PSST6* expression

#### Telencephalon

As indicate above for *PSST1*+ cells, *PSST6*+ cells are occasional or lack in the rostral pallium. More caudally, fairly abundant *PSST6*+ cells appear in medial, dorsal and lateral regions of the pallium (Figs. 1c-g, 4a-g). In the medial pallium, a dense population of intensely stained *PSST6*+ cells appears in a large area over the neuroporic recess (Fig. 4b-f). Caudally this median pallial population expands considerably and becomes continuous with paired condensations of intensely stained *PSST6*+ cells that are surrounded by less dense *PSST6*+ cells. No similar condensation of *PSST6*+ cells is seen in the dorsal and lateral regions of the pallium, which exhibit much lower density and fainter staining of *PSST6*+ cells. The olfactory bulbs show occasional *PSST6*+ cells (Fig. 1c).

**Figure 4.**
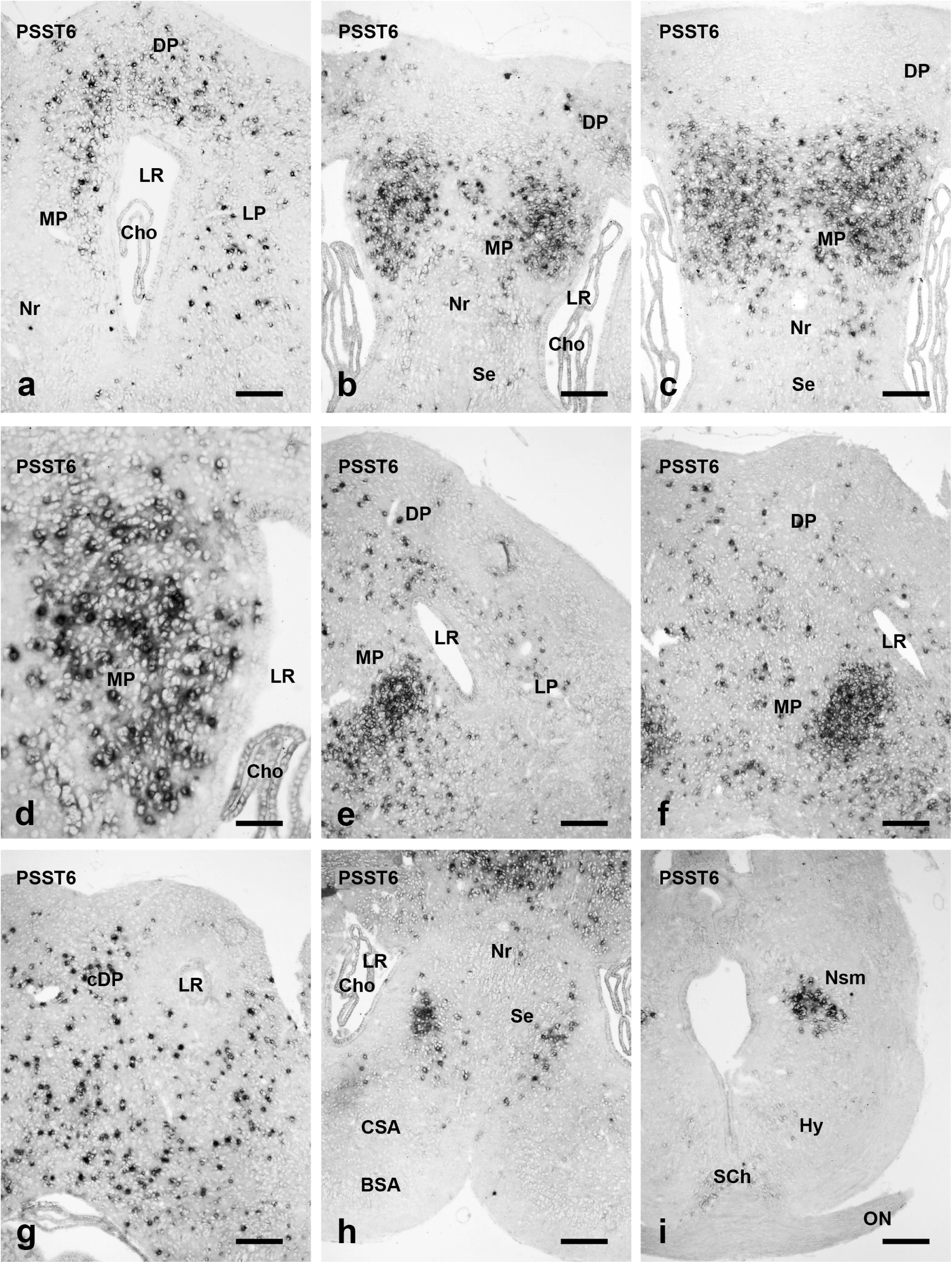
Low magnification photomicrographs of transverse brain sections of a juvenile catshark showing *PSST6* positive cells. These brain sections roughly correspond to the sections shown schematically at the right side in Figure 1. For abbreviations, see the list. Scale bars, 250 µm (a-c, e-i), 100 µm (d).

Double staining with *PSST6 in situ* hybridization and TH immunohistochemistry revealed that a proportion of caudal pallial *PSST6*+ cells were also TH-ir (Fig. 6a-c). In these cells blue *PSST6* positivity was distributed in the perikarya around the cell nucleus, whereas brown TH immunoreactivity was observed both in the perikaryon and cell processes. Colocalization with TH in the caudal pallium also indicates that these *PSST6*+/TH-ir neurons are different from *PSST1*+ pallial neurons of this region.

Scarce expression of *PSST6* was observed in the subpallium of juvenile catsharks. A small paired compact group of *PSST6*+ cells was observed in the medial septum (Fig. 4h), from which sparse cells extend caudolaterally. In the BSA, only occasional *PSST6*+ cells were observed (Fig. 1b-e), in sharp contrast with the distribution of *PSST1*+ cells. Subpallial *PSST6*+ cells did not show TH-ir immunoreactivity in double staining experiments. At the transition with the telencephalic peduncle, a compact ovoid group of small *PSST6*+ cells is observed forming a small protuberance close to the insertion of the anterior medullary velum (Figs. 1f-g, 4i). This group is in the nucleus of the stria medullaris (Fig. 6d) and ends close to the habenula. It may be considered part of the prethalamic eminence. The interstitial nucleus of the anterior commissure only shows few and pale *PSST6*+ cells (not shown).

#### Preoptic region and hypothalamus

The preoptic region of juvenile dogfish mostly lacks *PSST6*+ neurons, and only a few faintly stained neurons were observed in the lateral area at the level of the suprachiasmatic nucleus (Fig. 1h). A few double stained *PSST6*+/TH-ir cells were observed in this lateral preoptic/suprachiasmatic area among abundant TH-ir cells (Fig. 6e).

*PSST6+* neurons are scattered in areas dorsal to the lateral and posterior hypothalamic recesses that form part of the hypothalamic paraventricular organ (Fig. 1k-m), but positive cells are not present in this organ or in the median hypothalamic nucleus and lateral tuberal nucleus located over the median eminence. TH-ir cells were observed in some hypothalamic nuclei, sometimes aside or intermingled with *PSST6+* neurons (Fig. 6f-h), but no colocalization of TH with *PSST6* signal was observed.

#### Diencephalon

The main *PSST6*+ neuronal population observed in the dogfish diencephalon extends as a conspicuous cell band in intermediate medio-lateral levels of the prethalamus and posterior tubercle (Figs. 1h-k, 5a, 6f). In the posterior tubercle at the level of the nucleus of the saccus vasculosus (see Molist, Rodriguez-Moldes, & Anadon, 1992), *PSST6*+ cells form conspicuous groups lateral to it (Figs. 1m, 5b). Numerous prethalamic/posterior tubercular cells showed TH-ir immunoreactivity, but colocalization with *PSST6* was not observed in these cells. In the habenula, a small group of faint *PSST6*+ cells was observed near the lateral border (Fig. 1g), near the nucleus of the stria medullaris. Additionally, a few *PSST6*+ cells were observed in the nucleus of the medial longitudinal fascicle in periventricular location (Fig. 1j). These cells were not TH-ir.

**Figure 5.**
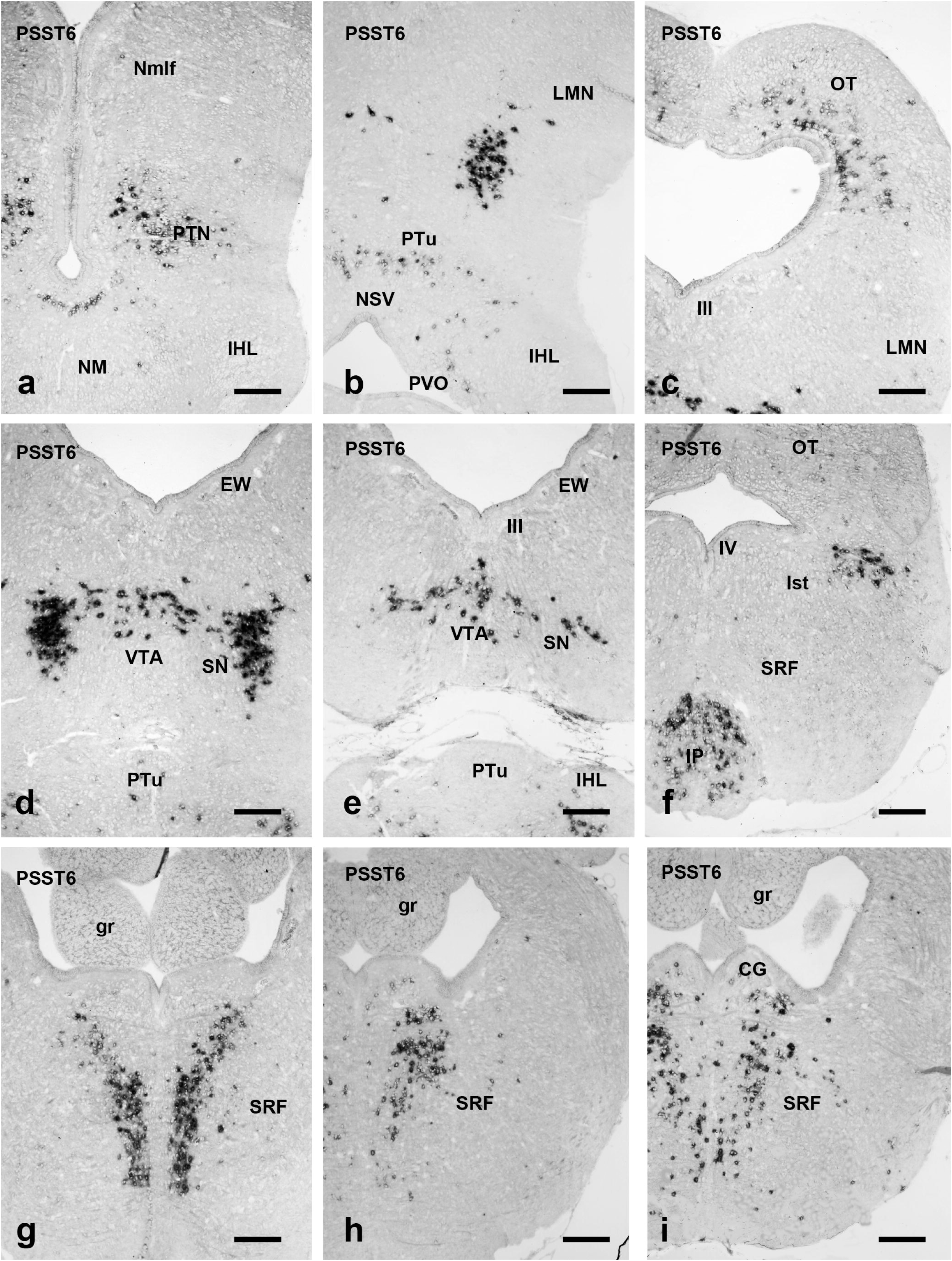
Continuation of Figure 4 showing low magnification photomicrographs of transverse brain sections with *PSST6* positive cells. These sections correspond to those shown schematically at the right side in Figure 1. For abbreviations, see the list. Scale bars, 250 µm.

#### Midbrain

In the catshark optic tectum, abundant *PSST6*+ neurons were observed in the stratum cellulare externum zona interna and in lateral regions of the stratum cellulare internum (Figs. 1i-n, 5c), which is in contrast with the distribution of *PSST1*+ cells mainly in the stratum cellulare externum zona externa (see above).

In the mesencephalic tegmentum a paired compact *PSST6*+ reticular nucleus is located in an intermediate region lateral to the oculomotor nerve, and more scattered *PSST6+* cells also extend between the two nuclei in the midline (Figs. 1-l-m, 5d-e). These reticular populations disappear at rostral isthmic levels caudally, and at the limit with the tegmental pretectal region rostrally. In double-stained sections for TH and *PSST6* (see above) these *PSST6*+ populations are co-distributed with the dopaminergic populations of the substantia nigra and ventral tegmental area, respectively (Fig. 6i). However, colocalization of *PSST6* and TH was not observed in any cell, unlike in the pallium and the preoptic/prethalamic populations. A similar SST-like-ir population of the midbrain tegmentum of the gummy shark was named red nucleus by Chiba, Honma, Ito, & Homma (1989).

**Figure 6.**
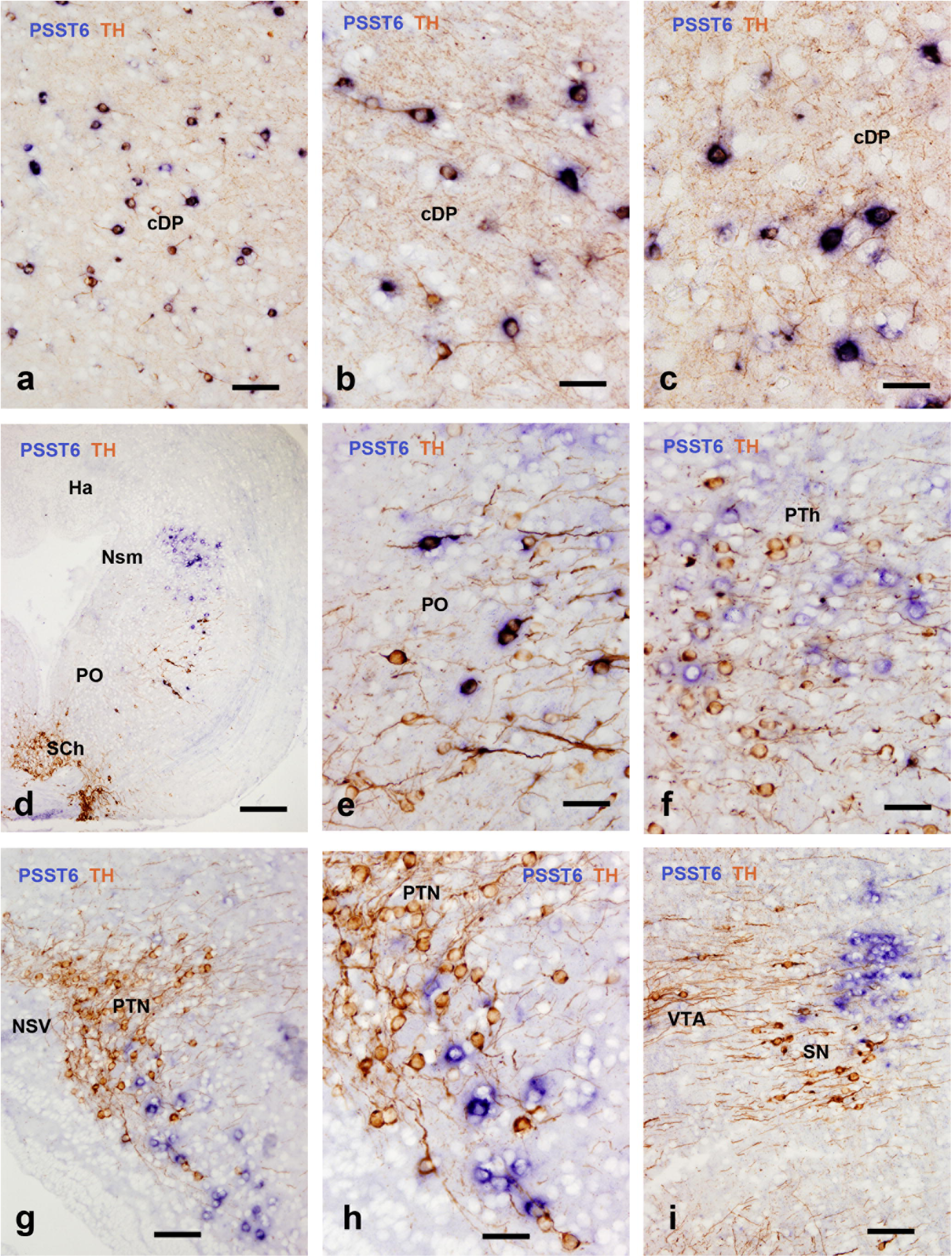
Photomicrographs of transverse brain sections of a juvenile catshark showing colocalization (**a-c**, **e**) or co-distribution (**d, f-i**) of *PSST6*+ and TH-ir structures. For abbreviations, see the list. Scale bars, 250 µm (d), 100 µm (a, g, i), 50 µm (b, c, e, f, h).

#### Rhombencephalon

The transition between the basal regions midbrain and the isthmus is marked by the appearance of the interpeduncular nucleus (isthmic) just caudal to the oculomotor nerve exit, and the end of the substantia nigra/ventral tegmental area populations. A rich population of *PSST6*+ neurons was observed in the dorsal and caudal regions of the interpeduncular nucleus, excepting the ventral interpeduncular neuropil (Figs. 1n-o, 5f). Just caudal to this nucleus, a paired band rich in *PSST6*+ cells of the superior reticular formation ascend dorsally parallel to the raphe and the medial longitudinal fascicle and reach a more massive population of *PSST6*+ cells located below the rhombencephalic central grey (Figs. 1p-r, 5g-h). Faint scattered *PSST6*+ cells are also observed ventral to the medial isthmic *PSST6*+ population. The *PSST6*+ medial reticular populations extend caudally in the trigeminal region (rhombomeres 2-3) till the level of entrance of the octaval nerve (Fig. 1s). At these levels, the central grey also contains a small population of *PSST6*+ neurons (Figs. 1r, 5i). A small lateral reticular population of *PSST6*+ cells that is separated from the meninges by the spino-cerebellar tract is also observed (Fig. 1q), but whether it is located in isthmic or trigeminal levels was not assessed. Caudally to the octaval nerve entrance, faintly stained *PSST6*+ cells were scattered in intermediate reticular areas (Fig. 1t). At the level of the vagal lobe, scarce *PSST6*+ reticular cells were observed in an area between this lobe, the caudal octavolateralis nucleus and the descending trigeminal nucleus (Fig. 1u). Near the obex, reticular *PSST6*+ cells were also observed below the commissural nucleus and spinal trigeminal nucleus, extending to the transition with the spinal cord (not shown). No *PSST6*+ cells were observed in the spinal cord.

### *PSST2* expression

Neurons expressing *PSST2* mRNA were observed in the median hypothalamic and lateral tuberal nuclei, the ventro-lateral hindbrain (Fig.7a-d) and the dorsal horn of the rostral spinal cord. Hypothalamic *PSST2*+ cells of the median hypothalamic nucleus extend from levels rostral to the origin of the lateral recesses of the third ventricle till the transition between the NM and the neurohypophysis (Figs. 7a-c, 8a-d). In rostral and intermediate levels positive cells are scattered near the midline between the third ventricle and the meninges (Fig. 8a-b). At levels in which the NM exhibits the shape of a paired bulge, numerous *PSST2+* cells appear grouped in the dense periventricular cell layer of the lateral tuberal nucleus, whereas the more ventral cells disappear gradually (Fig. 8c-d). The rostro-caudal extension of the periventricular population coincides with the extension of the adenohypophysis that lies just ventral to the bed of capillaries of the median eminence in juveniles.

In the caudal hindbrain there is a population of *PSST2*+ reticular neurons that extends rostro-caudally between the levels of the entrance of the glossopharyngeal nerve and a post-obecular level. These small cells form a small group located in a ventrolateral reticular area below the vagal sensory lobe (Figs. 7d, 8e-f).

At rostral spinal levels, some *PSST2*+ cells are scattered in the dorsal horn over the Stieda’s fasciculus medianus (direct vestibulo-spinal tract of elasmobranchs, which is located laterally to the central canal) or just below it. No positive cells located around the central canal were observed (not shown).

**Figure 7.**
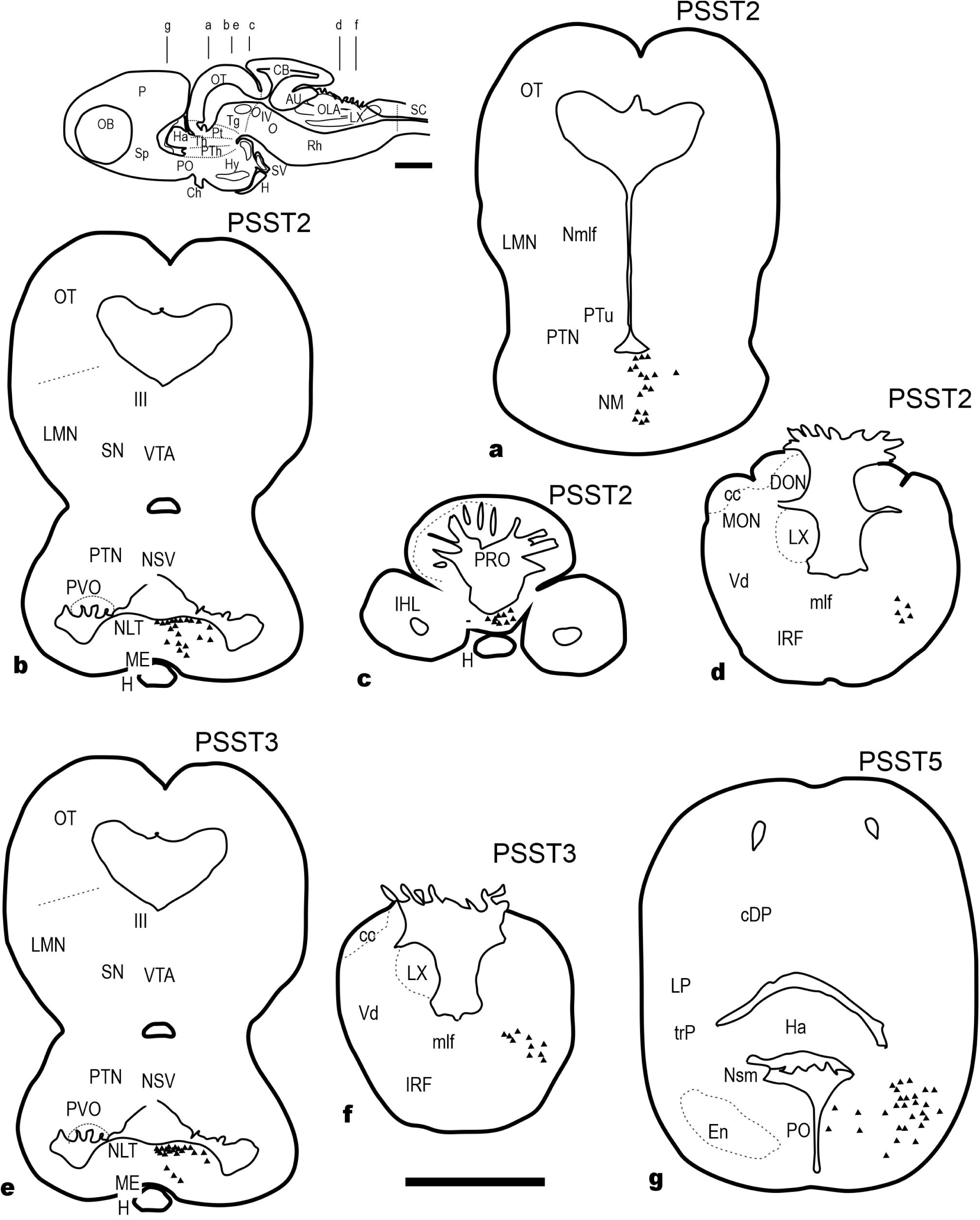
Schematic drawings of sections of juvenile catshark brains hybridized for *PSST2* (**a-d**), *PSST3* (**e-f**) and *PSST5* (**g**) showing the distribution of positive neurons (each black triangle represents a single cell in a 16µm-thick section). Anatomical structures are indicated in the left side of the left figurines. For abbreviations, see the list. The level of sections is indicated at the upper left corner in a schematic drawing of a juvenile brain. Scale bar: 1mm.

**Figure 8.**
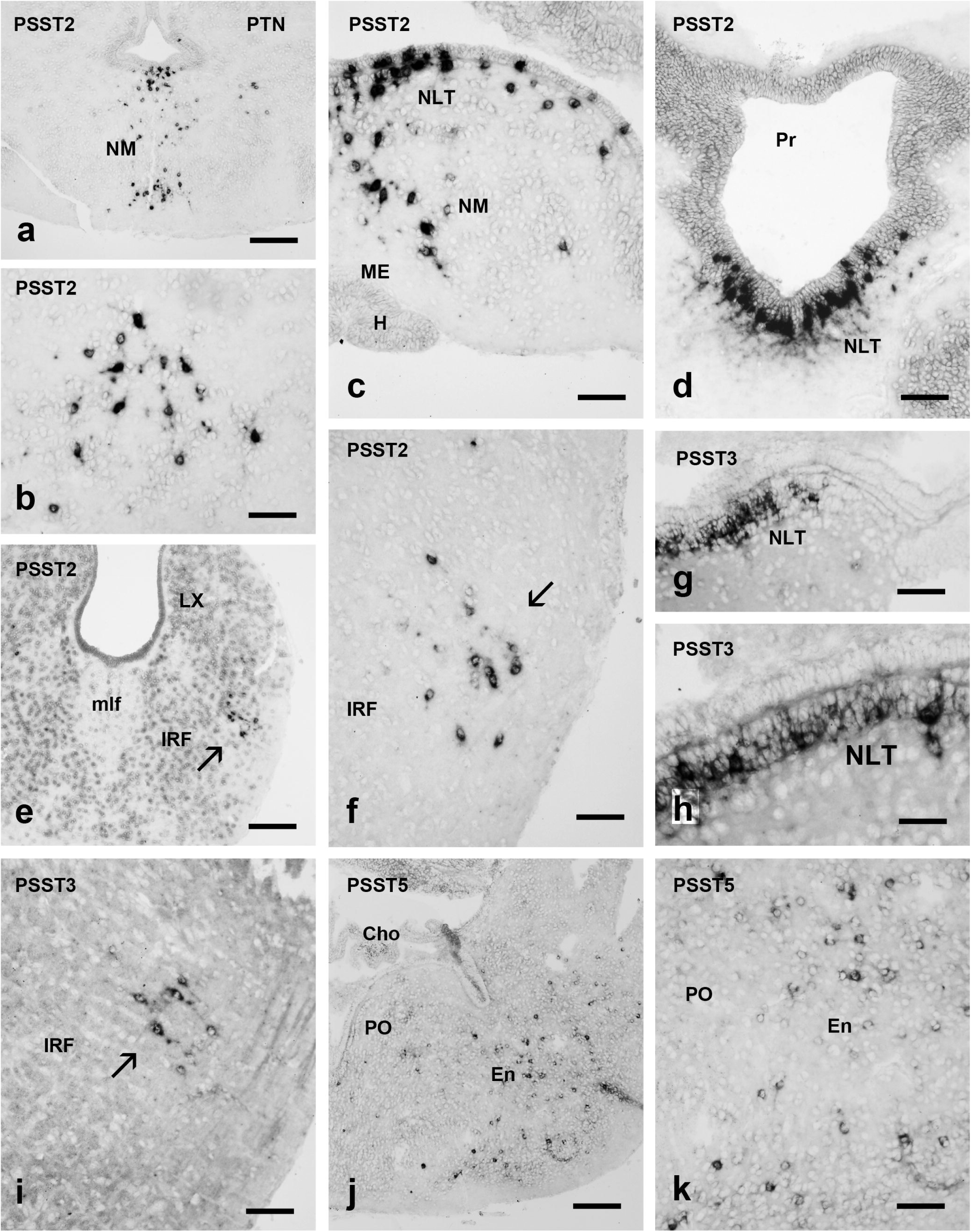
Photomicrographs of transverse brain sections of a juvenile catshark showing *PSST2* positive cells (**a-f**), *PSST3* positive cells (**g-i**), and *PSST5* positive cells (**j-k**). Arrows in e, f and i point to groups of lateral reticular neurons. These sections correspond to the brains shown schematically in Figure 7. For abbreviations, see the list. Scale bars, 250 µm (a, e, j), 100 µm (b, c, d, f, g, i, k), 50 µm (h).

### *PSST3* expression

Two *PSST3*-expressing neuronal populations were observed in the juvenile brain, one in the NM/NLT and the other in the caudal hindbrain (Figs. 7e-f), a pattern that reminds that reported above for the *PSST2*+ brain populations. However, no cells expressing this transcript were observed in the rostral spinal cord. Neurons of the NM/NLT are mostly located in the thick periventricular cell layer below the ependyma (Fig. 8g-h), but positive cells were also seen among the cells scattered within the neuropil below the dense periventricular layer. The NM/NLT lies over the plexuses of capillaries of the median eminence (Mellinger, 1964). The second *PSST3+* population is located near the meninges in the caudal hindbrain and consists of small reticular neurons in a ventrolateral region at the level of the glossopharyngeal nerve entrance (Fig. 7f, 8i). These small reticular cells have a bipolar appearance showing radial orientation.

### *PSST5* expression

In the transition between the impar telencephalon and the preoptic region there is a population with numerous *PSST5+* neurons that are scattered in a wide lateral region corresponding to the entopeduncular nucleus (Figs. 7g, 8j-k). Medially, only a few scattered *PSST5*+ cells were observed in the preoptic nucleus at the level of the preoptic recess. The magnocellular preoptic nucleus of catshark has been characterized immunohistochemically in this region (Meurling, Rodríguez, Peña, Grondona, & Pérez, (1996). The other brain regions of the juvenile catshark lack any positivity for the *PSST5* probe.

## DISCUSSION

### The five somatostatin genes exhibit different patterns of expression in the brain and spinal cord of catshark

Immunohistochemical results have revealed a widespread occurrence of SST-ir nerve cell bodies and fibers throughout the central nervous system of adult rats (Johansson, Hökfelt, & Elde, 1984) and other vertebrates, including sharks (Nozaki, Tsukahara, & Kobayashi, 1984; Chiba, Honma, Ito, & Homma, 1989). *In situ* hybridization studies reported a wide distribution of the mRNA expression of the single somatostatin gene found in mammals (Fitzpatrick-McElligott, Card, J.P., Lewis, M. E., & Baldino, 1988; Kiyama and Emson, 1990). The recent identification of five somatostatin genes in the catshark and other elasmobranchs revealed that the gene diversity of the somatostatin system in these fishes is similar to that previously reported in teleosts and close to that of the common gnathostome ancestor, but clearly different to that in mammals (Tostivint, Gaillard, Mazan, & Pézeron, 2019). Moreover, the gene synteny allowed studying the phylogeny of this gene family, in particular the homologies between paralogous genes. Thus, the *PSST3* and *PSST6* genes probably arose by tandem duplication of the *PSST1* and *PSST2* genes, respectively, although other alternatives are possible (see Tostivint, Gaillard, Mazan, & Pézeron, 2019). Comparison of patterns of expression of these five genes revealed by *in situ* hybridization in the catshark brain shows large differences between members of these genes. Thus, the wide brain distribution of *PSST1* and *PSST6* expression is in contrast with the limited expression of *PSST2*, *PSST3* and *PSST5*. Moreover, the distribution pattern of the two more widely expressed *PSST* genes in brain neurons (*PSST1* and *PSST6*) appears to be largely exclusive although double hybridization studies appear necessary to rule out the possibility of double expression in some neurons. The differential expressions reported here represent the first comprehensive study of the somatostatinergic systems in an elasmobranch.

Previous studies of elasmobranches using SST-like immunohistochemistry described the presence of somatostatinergic cells and/or fibers in various brain or spinal centers (Nozaki, Tsukahara, & Kobayashi, 1984; Chiba, Honma, Ito, & Homma, 1989; Cameron, Plenderleith, & Snow, 1990; Molist, Rodriguez-Moldes, & Anadon, 1992; Anadón, Molist, Pombal, Rodríguez-Moldes, & Rodicio, 1995; Alvarez-Otero, Perez, Rodriguez, & Anadón, 1996). SST-like-ir cell distribution reported in the gummy shark (Chiba, Honma, Ito, & Homma, 1989) reminds partially the distribution of *PSST1*+ cells observed here, although the number of cells observed was clearly lower and various populations were not noted. In these studies, the antibody used was raised against mammalian SST-14, whose aminoacidic sequence is identical to the putative mature SST peptides from catshark *PSST1* and *PSST5*. Future studies with antibodies specific for the other putative catshark somatostatins might also reveal differential distributions of some somatostatinergic fiber systems.

In the following we will discuss the patterns of expression of *PSSTs* in the catshark CNS in comparison with those reported in other vertebrates, whenever possible. In these comparisons, the reader should take in to account that the recent study by Tostivint, Gaillard, Mazan, & Pézeron (2019) suggest that the so-called *SST2* gene in chicken, lungfish, sturgeons and teleosts actually corresponds to *SST6*. For convenience, in the text we will keep the nomenclature used in original studies of each species.

### Wide but differential expression of PSST1 and PSST6 genes in pallial interneurons

SST is expressed in subsets of GABAergic interneurons of the mammalian pallium (hippocampus, cortex) and appears to play a key role in their inhibitory circuits (Urban-Ciecko and Barth, 2016; Adler, Zhao, Shin, Yasuda, & Gan, 2019). Moreover, a number of subpopulations of SST neurons can be distinguished by morphology, connectivity, laminar location, firing properties, and expression of molecular markers (Yavorska and Wehr, 2016). Interestingly, wide expression of *PSST1* and *PSST6* genes is also observed in the catshark pallium, forming at least two sets of neurons. The catshark pallium presents a poor laminar organization, but exhibits numerous morphological cell types judging from studies with the Golgi method (Manso and Anadón, 1993), including areas with primitive pyramidal neurons and other large neurons. The scattered distribution and small size of *PSST1*+ and *PSST6*+ pallial neurons may suggest that they are local interneurons, but this has not been assessed. The presence of SST-like-ir pallial neurons was reported in the gummy shark using immunohistochemistry (Chiba, Honma, Ito, & Homma, 1989). In mammals, all pallial somatostatinergic neurons represent subtypes of GABAergic interneurons (Esclapez and Houser, 1995), and thus they probably originate from the embryonic subpallial ganglionic eminences (Anderson, Marín, Horn, Jennings, & Rubenstein, 2001). Numerous GABAergic cells are found in the catshark pallium and they also appear to have a subpallial origin, migrating secondarily to the pallium (Carrera, Ferreiro-Galve, Sueiro, Anadón, & Rodríguez-Moldes, 2008b; Quintana-Urzainqui, Rodríguez-Moldes, Mazan, & Candal, 2015). Although these parallelisms with mammals are suggestive of colocalization of SST and GABA in cells of the pallium, if *PSST1*+ and/or *PSST6*+ neurons of the catshark pallium are also GABAergic need be investigated. SST-ir neurons and large number of fibers have been reported in the pallium (dorsal telencephalic area) of ray-finned fishes (trout: Becerra, Manso, Rodríguez-Moldes, & Anadón, 1995; sturgeon: Adrio, Anadón, & Rodríguez-Moldes, 2008). Moreover, mRNA expression of the *PSST1* gene was also reported in scattered neurons of the pallium of sturgeon (Trabucchi et al., 2002) and goldfish (Canosa, Cerdá-Reverter, & Peter, 2004). Unlike the catshark *PSST6*, the other *PSST* genes of these ray-finned fishes show no expression in the pallium. However, expression of both *PSST1* and *PSST2* was reported in the frog pallium (Tostivint et al., 1996). Together, these findings suggest partially conserved patterns in the somatostatinergic pallial system of jawed fishes. A loose network of SST-ir fibers in the pallial neuropil was noted in the gummy shark (Chiba, Honma, Ito, & Homma, 1989), although the origin of fibers was not established. Immunohistochemical data in the trout reveal that the pallium was the brain region most richly innervated by SST-ir fibers, suggesting a great importance of somatostatin in pallial circuits (Becerra, Manso, Rodríguez-Moldes, & Anadón, 1995). The presence of two types of somatostatinergic pallial neurons adds neurochemical complexity to the cellular diversity of the catshark pallium, which also has interneurons expressing choline acetyltransferase (Anadón et al., 2000), GABA (Carrera, Ferreiro-Galve, Sueiro, Anadón, & Rodríguez-Moldes, 2008b), glycine (Anadón, Rodríguez-Moldes, & Adrio, 2013), tyrosine hydroxylase (Carrera et al., 2005; Carrera, Anadón, & Rodríguez-Moldes, 2012), thyrotropin-releasing hormone (Teijido, Manso, & Anadón, 2002) and met-enkephalin (Quintana-Urzainqui et al., 2012). Here, colocalization between TH and *PSST6* is reported in pallial neurons (see below), but if any *PSSTs* are colocalized with the other substances was not investigated.

### Preferential expression of PSST6 in the catshark subpallium

The adult elasmobranch subpallium shows a cellular organization that is difficult to compare with that of lampreys and bony fishes. Following Smeets, Nieuwenhuys, & Roberts (1983) the subpallium is composed of an extensive dense cellular lamina (BSA) that is separated from the ventricle by a wide central basal area and a thin ventral periventricular area. Medially, a septal region joins the pallium dorsally at the level of the neuroporic recess and ventrolaterally to the BSA. Between the BSA and the retrobulbar region there is an ill-organized cellular region considered as the striatum by Smeets, Nieuwenhuys, & Roberts (1983). Toward the telencephalic peduncle (at the level of the anterior commissure) this highly developed basal region is substituted by the poorly organized entopeduncular area laterally and the interstitial nucleus of the anterior commissure medially. New views of the subpallium have emerged from genoarchitectonic studies of the embryonic subpallium (reviewed in Rodríguez-Moldes, Santos-Durán, Pose-Méndez, Quintana-Urzainqui, & Candal, 2017). In catshark embryos, the subpallium primordium was characterized by the early expression of GABAergic markers (GAD, glutamic acid decarboxylase) and *Dlx2* (Carrera, Ferreiro-Galve, Sueiro, Anadón, & Rodríguez-Moldes, 2008b; Ferreiro-Galve et al., 2008; Quintana-Urzainqui et al., 2012; Quintana-Urzainqui, Rodríguez-Moldes, Mazan, & Candal, 2015). The medial region of the subpallium expresses *Nkx2.1* in a band comprising the midline and immediately adjacent lateral territories, and sonic hedgehog (shh) in the midline. The embryonic primordium expressing these early markers was proposed to correspond with the embryonic median ganglionic eminence of tetrapods that originates the pallidum. The pattern of gene expression observed in the remainder subpallial primordium corresponds with the mammalian lateral ganglionic eminence that originates the striatum, which does not express *Nkx2.1* and shh, as also shown in catshark (Quintana-Urzainqui et al., 2012). According to this embryo-centered point of view, the catshark subpallium is mainly comprised of striatum and pallidum, and perhaps a septal region rostrally (Quintana-Urzainqui et al., 2012; Quintana-Urzainqui, Rodríguez-Moldes, Mazan, & Candal, 2015). These authors also demonstrate complex cell migratory routes in the embryonic telencephalon from the subpallium to the pallium and to the olfactory bulbs, which led to large transformations after the embryonic stage 32. Unresolved questions remain about the actual correspondence between subpallial and pallial territories of the adult catshark with their amniote counterparts.

The presence of SST-ir cells in the septum and basal area was reported in the gummy shark (Chiba, Honma, Ito, & Homma, 1989). With regard to the expression of *PSST* genes in the catshark subpallium, *PSST1* was mainly expressed in cells of the basal and central superficial areas, whereas *PSST6* and the other *PSSTs* are not expressed there. Both *PSST1* and *PSST6* are expressed in the septum of Smeets, Nieuwenhuys, & Roberts (1983), although with different patterns. *PSST1* is poorly expressed in cells of the striatum of these authors and lacks in the interstitial nucleus of the anterior commissure, which shows few *PSST6*+ cells. In ray-finned fishes as sturgeon, *PSST1* and *PSST2* (now renamed *PSST6* according to Tostivint, Gaillard, Mazan, & Pézeron, 2019) genes are expressed in the subpallium (ventral telencephalic area) although with different patterns (Trabucchi et al., 2002), but it is not possible to establish equivalences with the expression patterns observed in catshark. Two somatostatin genes are abundantly expressed in the lateral region of the goldfish telencephalic ventral area (Canosa, Cerdá-Reverter, & Peter, 2004). Somatostatin-like immunochemistry reveals positive cells in the sturgeon subpallium, mainly in the lateral region of the ventral telencephalic area (Adrio, Anadón, & Rodríguez-Moldes, 2008).

### Olfactory bulbs

The catshark olfactory bulbs show scarce *PSST1*+ cells in the inner granular layer and lack *PSST6* expression. An intriguing result is the small group of intensely *PSST1*+ cells found at the level of the glomerular layer. Unlike the catshark, *PSST-III* was intensely expressed in the internal cell layer (ICL) of the goldfish olfactory bulb (Canosa, Cerdá-Reverter, & Peter, 2004), whereas in sturgeon the olfactory bulbs do not express *PSST1* or *PSST2* genes (Trabucchi et al., 2002).

### PSST1 is specifically expressed by CSF-contacting neurons of the paraventricular/posterior recess organ and central canal

The paraventricular and posterior recess organs have been characterized in catshark and other elasmobranches with histochemistry, immunohistochemistry and electron microscopy (Wilson and Dodd, 1973; Rodríguez-Moldes, 1986; Rodríguez-Moldes and Anadón, 1987; Meredith and Smeets, 1987; Meurling and Rodríguez, 1990; Rodríguez-Moldes et al., 1993; Molist, Rodríguez-Moldes, & Anadón, 1993; Carrera, Molist, Anadón, & Rodríguez-Moldes, 2008a; Anadón, Rodríguez-Moldes, & Adrio, 2013): these organs exhibit numerous dopaminergic, serotonergic and glycinergic CSF-c neurons. In *Squalus acanthias* and *Raja radiata* they showed numerous somatostatinergic CSF-c neurons showing colocalization with serotonin (Meurling and Rodríguez, 1990). Our *in situ* hybridization results reveal strong expression of *PSST1* mRNA in CSF-contacting neurons of both the hypothalamic paraventricular and posterior recess organs, and the spinal cord around the central canal. On the other hand, the other four *PSST* genes were not expressed in these organs. These *in situ* hybridization results are in agreement with the immunohistochemical observations of distribution of SST-like expression in CSF-c cells of these organs by Chiba, Honma, Ito, & Honma (1989) and Meurling and Rodríguez (1990). SST-like-ir CSF-c populations are also present in the lamprey hypothalamus (Wright, 1986; Yáñez, Rodríguez-Moldes, & Anadón, 1992).

With regard the catshark spinal cord, *PSST1*+ CSF-c cells show a horseshoe distribution around dorsal and lateral regions of the central canal, lacking in its ventral part. This distribution is complementary to that the spinal TH-ir CSF-c cells, which are located in the ventral region of the central canal (Sueiro, Carrera, Rodríguez-Moldes, Molist, & Anadón, 2003). This was confirmed here with a combination of *in situ* hybridization and immunostaining. Judging from the distribution of GABAergic CSF-c cells all around the catshark central canal, and the colocalization of TH and GAD in ventral CSF-c neurons (Sueiro, Carrera, Molist, Rodríguez-Moldes, & Anadón, 2004), this strongly suggests the *PSST1*+ cells are also GABAergic. Dorsal CSF-c cells, but not ventral cells, also express glycine immunoreactivity (Anadón, Rodríguez-Moldes, & Adrio, 2013), which might colocalize with *PSST1*. Presence of SST and colocalization with GABA immunoreactivity in spinal CSF-c cells was already reported in lampreys (Buchanan, Brodin, Hökfelt, Van Dongen, & Grillner, 1987; Christenson, Alford, Grillner, & Hökfelt, 1991; Jalalvand, Robertson, Wallén, Hill, & Grillner, 2014). In the catshark these spinal cells appear to give rise to the somatostatinergic fibers innervating the glomerular fields of the spinal marginal nucleus (Anadón, Molist, Pombal, Rodríguez-Moldes, & Rodicio, 1995). This catshark marginal nucleus also exhibits glycinergic neurons (Anadón, Rodríguez-Moldes, & Adrio, 2013), and similar glycinergic edge cells were also found in lampreys (Villar-Cerviño, Holstein, Martinelli, Anadón, & Rodicio, 2008), revealing a conserved pattern in these basal vertebrate groups. In lampreys, these spinal CSF-c cells modulate the edge cell activity in response to stretching of the spinal cord during swimming (Grillner, Williams, & Lagerbäck, 1984; Jalalvand, Robertson, Wallén, Hill, & Grillner, 2014). Recent studies in hypothalamic and central canal CSF-c cells of lampreys and zebrafish reveal that these CSF-c neurons are chemo- and mechanoreceptive, sensing pH changes and modulating the locomotor activity (Böhm et al., 2016; Jalalvand et al., 2018).

Available data in ray-finned fishes reveal that both *PSST1* and *PSST2* genes are expressed in the hypothalamic nucleus of the periventricular organ of sturgeon (Trabucchi et al., 2002). Some somatostatinergic CSF-c neuronal populations were described in sturgeon with SST immunohistochemistry, including the preoptic recess organ, the anterior tuberal nucleus, the posterior recess organ, and the spinal central canal (Adrio, Anadón, & Rodríguez-Moldes, 2008), which is roughly similar to the distribution observed in catshark with the sum of hybridization results with the *PSST1* (PVO, CC) and *PSST2* (lateral tuberal nucleus, which possibly corresponds to the sturgeon anterior tuberal nucleus) genes.

### Possible hypophysiotropic somatostatinergic hypothalamic populations

The structure of the median eminence of the catshark and its vascularization has been thoroughly described by Mellinger (1964). This eminence is located below the medial hypothalamus and receives fibers from neurosecretory cells (Knowles, 1965). Some of the axons projecting to the pituitary in teleost originate from ventromedial hypothalamic populations (Johnston and Maler, 1992; Anglade, Zandbergen, & Kah, 1993; Holmqvist and Ekström, 1995). As far as we know the neurons afferent to the median eminence in elasmobranchs have not been identified experimentally. However, abundant SST-like-ir fibers coursed in the hypophyseal median eminence and neurointermediate lobe of the gummy shark (Chiba, Honma, Ito, & Homma, 1989). The lateral tuberal nucleus and the median hypothalamic nucleus of the catshark showed neurons expressing *PSST1* (MHN), *PSST2* (NLT and MHN) and *PSST3* (MHN), but not cells expressing *PSST5* or *PSST6*. The MHN and LTN are located close to the median eminence, and *PSST*+ cells in this region might send fibers to it, which thus might control the secretion of growth hormone or other pituitary hormones. Further studies are needed to reveal the hypophysiotrophic populations of the catshark hypothalamus.

A population of *PSST6*+ cells was observed in the posterior tubercle lateral to the nucleus of the saccus vasculosus (NSV), a region which shows the presence of somatostatin-ir cells in catshark (Molist, Rodriguez-Moldes, & Anadon, 1992). The NSV receives a heavy neural projection from the hypothalamic saccus vasculosus, a neuro-ependymo-vascular structure characteristic of elasmobranchs and many ray-finned fishes (see Yáñez et al., 1997; Sueiro et al., 2007). However, the NSV proper only receives a few SST-like-ir fibers, probably not originated from the saccus vasculosus since no SST-like-ir neurons (Chiba, Honma, Ito, & Homma, 1989) or *PSST* expression (present work) was observed in this structure.

### Banded expression of somatostatin in the diencephalon

The main *PSST1*+ populations of the diencephalon appear to show a segmental-like distribution in the form of dorso-ventral bands in the prethalamus, thalamus and pretectum, suggesting that there is a main band in each prosomere. The main *PSST6*+ neuronal population observed in the catshark diencephalon also extends in a band between the prethalamus and posterior tubercle. In goldfish, *PSST1*, *PSST2* and *PSST3* mRNAs are expressed in nuclei known to pertain to the three diencephalic prosomeres (Canosa, Cerdá-Reverter, & Peter, 2004), but the pattern appears more complex than in catshark and its segmental organization was not analyzed. *PSST1* and *PSST2* expression was also noted in the prethalamus (ventral thalamus) and thalamus (dorsal thalamus) of sturgeon (Trabucchi et al., 2002).

### Upper and lower layers of the optic tectum differentially express PSST1 and PSST6

Experimental studies on the primary visual pathways in catshark reported that optic tectum receives a major retinal projection on external tectal layers (Smeets, 1981; Repérant, Miceli, Rio, Peyrichoux, Pierre, & Kirpitchnikova, 1986), as in most vertebrates. The catshark optic tectum also receives afferents from various brain nuclei and regions extending from the pallium to the spinal cord (Smeets, 1982), probably reaching deeper tectal layers. Notably, in the catshark optic tectum a complementary pattern of expression of *PSST1* and *PSST6* genes was observed. *PSST1*+ cells are mainly distributed through the stratum cellulare externum zona externa, whereas *PSST6*+ cells are distributed in the stratum cellulare externum zona interna and in lateral regions of the stratum cellulare internum. This may suggest specialization of outer *PSST1*+ cells for visual circuits and inner *PSST6*+ cells for other sensory and non-sensory afferents, respectively. Several types of neurons have been reported with Golgi methods in the catshark optic tectum, and some of them exhibit radial dendrites coursing through two of more layers (Manso and Anadón, 1991a), but ascription of *PSST+* cells to these cell types is not possible. The stratum cellulare externum of the catshark optic tectum is synaptically complex, exhibiting presynaptic dendrites and other unconventional synapses (Manso and Anadón, 1991b). In the goldfish optic tectum, the cells expressing the *PSST1* and *PSST3* genes are both distributed in the periventricular gray zone (Canosa, Cerdá-Reverter, & Peter, 2004), whereas in sturgeon tectum only the *PSST2* gene is expressed (Trabucchi et al., 2002). Expression of *PSST1,* but not of *PSST2,* was reported in the frog optic tectum (Tostivint et al., 1996). In the adult rat, *PSST* mRNA is expressed in numerous superior collicular neurons showing zonal distribution (Harvey, Heavens, Yellachich, & Sirinathsinghji, 2001). *PSST* mRNA expressing cells were rarely found in the stratum zonale (SZ) or upper stratum griseum superficiale (SGS), but formed a prominent tier in the lower third of SGS and in the upper stratum opticum (SO), a distribution that reminds the sum of that observed in catshark with two different *PSST* genes. Thus, specialization of SST in different tectal subcircuits depends on differential gene expression in sharks but not in mammals.

### Tegmental expression of PSST genes and its topographical relation with the catecholaminergic substantia nigra/ventral tegmental area

In the midbrain tegmentum of catshark, the most conspicuous *PSST1*+ cell population was observed in the lateral reticular area whereas a paired compact *PSST6*+ reticular nucleus is located in an intermediate lateral region and more scattered *PSST6*+ cells extend in the midline. Faint *PSST1*+ cells were also observed intermingled with TH-ir neurons of the ventral tegmental area and the substantia nigra, but without forming any conspicuous group as *PSST6*+ cells. The presence of a group of large SST-like-ir neurons was reported in the tegmentum of the gummy shark and ascribed to the red nucleus (Chiba, Honma, Ito, & Homma, 1989). By its position and cell size it may correspond with the main *PSST6*+ population observed in the catshark midbrain tegmentum. The lateral *PSST1+* population may be part of the region known as the lateral mesencephalic nucleus that also contains a population of glycinergic cells (Anadón, Rodríguez-Moldes, & Adrio, 2013). In the spiny dogfish, probably also in other elasmobranchs, the lateral mesencephalic nucleus receives secondary octavolateral projections (Boord and Northcutt, 1988). This caudal midbrain *PSST1*+ population is located in a characteristic lateroventral outgrowth that contains a conspicuous precerebellar population (Pose-Méndez, Candal, Adrio, & Rodríguez-Moldes, 2014), and that may correspond to the nucleus H of guitarfish (Fiebig, 1988). Although these catshark precerebellar neurons were considered rhombencephalic, it is probable that they are mesencephalic judging from the location of *PSST1+* cells in relation to midbrain TH-ir populations. Prominent precerebellar cell populations are found in the midbrain tegmentum of sturgeon (Huesa, Anadón, & Yáñez, 2003) and in the lateral valvular nucleus of teleosts (see Folgueira, Anadón, & Yáñez, 2006), but not in the rostral rhombencephalic tegmentum. If the catshark tegmental *PSST1+* cells project to the cerebellum was not assessed. This suggests that somatostatinergic tegmental populations appear to be differentially involved in different neural circuits. In the frog and sturgeon midbrain tegmentum, both *PSST1* and *PSST2* genes are expressed (Tostivint et al., 1996; Trabucchi et al., 2002), but equivalences between catshark nuclei and those of these species are not defined.

### Expression of PSSTs in the catshark habenulo-interpeduncular system

Conspicuous expression of *PSST6*, but not of the other four *PSSTs*, is observed in numerous cells of the catshark interpeduncular nucleus (IP). The interpeduncular nucleus of the leopard shark and the gummy shark contains SST-like-ir neurons (Nozaki, Tsukahara, & Kobayashi, 1984; Chiba, Honma, Ito, & Homma, 1989), which may correspond to the *PSST6*+ group reported here. Glycinergic neurons were also reported in similar location in the catshark IP (Anadón, Rodríguez-Moldes, & Adrio, 2013), but if there is colocalization with *PSST6* was not investigated. In sharks, as in other vertebrates, the IP receives a heavy cholinergic habenular projection via the fasciculus retroflexus (Anadón et al., 2000; Giuliani, Minelli, Quaglia, & Villani, 2002). Two somatostatin genes are expressed in the IP of goldfish (*PSST1* and *PSST3;* Canosa, Cerdá-Reverter, & Peter, 2004) and frog (*PSST1* and *PSST2;* Tostivint et al., 1996). Moreover, SST-14-ir cells are found in a specific IP subnucleus in mice (Quina et al., 2017). The mammalian IP is the target of the medial habenular nucleus via the fasciculus retroflexus, and in turn projects to several brain regions, as shown in detail in mice (Quina, Harris, Zeng, & Turner, 2017). With regard the habenula, a small group of *PSST6*+ cells was found in the catshark habenula but no similar SST-like-ir cells were reported in the gummy shark (Chiba, Honma, Ito, & Homma, 1989). Cells expressing *PSST1* were reported in the goldfish habenula (Canosa, Cerdá-Reverter, & Peter, 2004).

To our knowledge, no *PSST*-expressing habenular cells were reported in other vertebrates. Some afferents to the habenula of the Arabian bamboo shark (*Chiloscyllium arabicum*) have been traced experimentally (Giuliani, Minelli, Quaglia, & Villani, 2002); these include the BSA, the entopeduncular nucleus, the posterior tubercle, thalamus and septum. Interestingly, the entopeduncular nucleus is the only region expressing *PSST5* in the catshark. This region also shows scarce *PSST1*+ cells. In teleosts, the habenula also receives fibers from the telencephalon (olfactory bulb, subpallium and entopeduncular nucleus), posterior tubercle, hypothalamus and raphe, the entopeduncular nucleus representing the main afferent nucleus (Yáñez and Anadón, 1996; Turner et al., 2016). In goldfish, the entopeduncular nucleus shows cells expressing *PSST3* and *PSST1* (Canosa, Cerdá-Reverter, & Peter, 2004), but both genes have a wide pattern of expression in the brain, which is unlike the catshark *PSST5*. The habenula is a highly conserved nucleus that forms part of the limbic system and is involved in a number of functions (Hsu et al., 2014).

### PSST and the viscerosensory lobe

In the catshark vagal lobe numerous *PSST1*+ small cells are observed in its dorsal half intermingled with TH-ir neurons, which form a separate population. In the catshark, a population of *PSST1*+ reticular cells also accompanies the solitary tract-vagal lobe at the level of the sulcus limitans, laterally to the motor nucleus. The other *PSSTs* are not expresses here. In the gummy shark (Chiba, Honma, Ito, & Homma, 1989), the vagal lobe shows numerous SST-like-ir fibers and some cells. In the vagal lobe of sturgeon there is also expression of *PSST1* but not *PSST2* (Trabucchi et al., 2002), but distribution of SST-ir cells is not parceled as in the catshark (Adrio, Anadón, & Rodríguez-Moldes, 2008). The goldfish has a complex viscerosensory region with three lobes (facial, glossopharyngeal and vagal), the vagal lobe being formed by several sensory and motor layers (Morita and Finger, 1985; Finger, 2008). *In situ* hybridization studies in goldfish (Canosa, Cerdá-Reverter, & Peter, 2004) indicate that *PSST1+* cells are distributed in superficial (sensory) layers, whereas *PSST3* is expressed in cells of deep (motor) layers. In rat, SST expression was also noted in the homologous nucleus of the solitary tract using ^32^P-labeled oligonucleotide probes (Priestley, Réthelyi, M., & Lund, 1991), indicating conserved patterns of expression. In mice, the solitary nucleus shows a few GABAergic are also SST-ir (Wang and Bradley, 2010), suggesting that somatostatinergic cells of this nucleus are inhibitory interneurons, as those found in the cortex. If this also occurs in the viscerosensory lobes of catshark and other vertebrates needs to be investigated.

### The hindbrain reticular formation and expression of PSSTs

The hindbrain reticular region *sensu amplio* is formed of scattered neurons and neuropil intermingled with a large number of longitudinal and transverse fiber fascicles. These areas are highly developed in the catshark hindbrain (Smeets, Nieuwenhuys, & Roberts, 1983), occupying most of the basal plate-derived region excepting the areas occupied by defined nuclei as the motor nuclei (Anadón et al., 2000; Rodríguez-Moldes et al., 2011), or the inferior olive. The hindbrain reticular formation has been subdivided in isthmic, superior, medius and inferior reticular nuclei mainly attending to the distribution of large reticular cells (see Smeets, Nieuwenhuys, & Roberts, 1983). Reticulospinal cells are the origin of major descending pathways to the spinal cord and were found in all these parts of the reticular formation (Smeets and Timerick, 1981; Timerick, Roberts, & Paul, 1992). Most of the reticulospinal neurons reported by these authors were large neurons, whereas in general somatostatinergic reticular neurons are small or medium-sized cells often distributed in lateral and/or medial groups, as noted above in the midbrain reticular region. The distribution of *PSST*+ reticular populations might be described using transverse segmental (rhombomeres) and longitudinal columnar patterns (Rodríguez-Moldes et al., 2011). A clear segmental pattern of reticulospinal cells has been observed in cyprinids (Metcalfe, Mendelson, & Kimmel, 1986; Lee, Eaton, & Zottoli, 1993; Gilland, Williams, & Lagerbäck, 2014) although not in sharks. With regards the hindbrain columnar organization, recent studies in the larval zebrafish hindbrain revealed a longitudinal patterning in which neurochemically different (glutamatergic, GABAergic, glycinergic) neuron stripes interleave in transverse sections (Higashijima, Mandel, & Fetcho, 2004; Kinkhabwala et al., 2011). Although these stripes become deeply modified during development, similar glycinergic populations may be observed in adult zebrafish (Barreiro-Iglesias et al., 2013). Moreover, reticular populations may be located in peripheral (superficial), intermediate or periventricular location, which probably reflects the neuronal birth time (early, intermediate or late) and/or the existence of radial and/or tangential migrations as those described in catshark hindbrain (Carrera, Molist, Anadón, & Rodríguez-Moldes, 2008a; Pose-Méndez, Candal, Adrio, & Rodríguez-Moldes, 2014).

The main groups of *PSST*-expressing reticular cells observed in the catshark rhombencephalon can be defined by a combination of these patterning criteria. Examples of *PSST1+* populations are the strongly *PSST1*+ reticular cells in a mediolateral region at the level of the interpeduncular nucleus (rhombomere 1), the groups of small *PSST1*+ cells observed extending in a vertical band near the midline between the central grey and the ventral surface, or cells migrated laterally forming a band parallel to the meninges. The strong *PSST1*+ reticular cells accompanying the extension of the solitary tract-vagal lobe are a lateral population. Some examples of *PSST6+* populations are the paired cell band near the medial longitudinal fascicle in the isthmic tegmentum, or the *PSST6*+ medial reticular population that extends in rhombomeres 2-3. Finally, *PSST2+* and *PSST3+* reticular neurons populations located near the ventrolateral meninges may be considered to form part of the far-migrated lateral reticular nucleus (LRN).

### The cerebellar system

The cellular organization of the catshark cerebellum has been characterized with microscopic and immunohistochemical methods (see Alvarez-Otero, Pérez, Rodríguez, Adrio, & Anadón, 1995; Anadón et al., 2009, 2013). No positive cells for any of the five *PSSTs* were observed in the catshark cerebellum with ISH (not shown) or in the gummy shark cerebellum with SST-like immunohistochemistry (Chiba, Honma, Ito, & Homma, 1989). Unlike in the shark cerebellum, SST-ir cells (Golgi cells, some Purkinje cells) were described in mammals with immunohistochemistry (Vincent et al., 1985) and *in situ* hybridization (Inagaki et al., 1989). Tracing experiments in catshark indicate that cells of the LRN and other regions containing *PSST+* cells project to the cerebellum (Pose-Méndez, Candal, Adrio, & Rodríguez-Moldes, 2014). Moreover, SST-ir fibers were observed in the cerebellar nucleus of catshark (Alvarez-Otero, Perez, Rodriguez, & Anadón, 1996; Anadón et al., 2009), but not in the cerebellar cortex of the gummy shark (Chiba, Honma, Ito, & Homma, 1989). In mice, evidence of innervation of the cerebellar granular layer by SST-28 immunoreactive mossy fibers has been presented (Armstrong et al., 2009). If *PSSTs* are expressed in some of the precerebellar cells reported in catshark (Pose-Méndez, Candal, Adrio, & Rodríguez-Moldes, 2014) need be investigated.

### Colocalization of TH immunoreactivity and PSST6 mRNA expression in brain neurons

Initially, we decided to perform double staining with TH immunohistochemistry in catshark as a marker to clarify the topography of some *PSST1*+ or *PSST6*+ neuronal populations, because the expression of this enzyme in several brain regions containing somatostatinergic neurons is well characterized (Carrera et al., 2005; Carrera, Anadón, & Rodríguez-Moldes, 2012). Whereas most of these populations did not show colocalization of *PSSTs* and TH, to our surprise two populations in the pallium and in the lateral preoptic/suprachiasmatic area showed double-labeled cells for TH-ir and *PSST6+*. The presence of TH-ir cortical neurons has been previously reported in the pallium of mammals, elasmobranchs and some reptiles (see Smeets and González, 2000). In the rat, double immunohistochemical studies indicate that very few TH-ir cortical neurons contained immunoreactivity for SST (Asmus et al., 2007), which is similar to our observations in catshark with TH and the *PSST1* probe. However, numerous catshark pallial neurons exhibit both *PSST6* positivity and TH immunoreactivity, indicating for the first time in a vertebrate the wide coexistence of a catecholamine-synthesizing enzyme and the expression of one SST gene in pallial cells. It is difficult to say if it might represent an ancestral feature of jawed vertebrates that was lost in evolution, because there are no TH-ir neurons in the pallium of most non-mammalian vertebrates (Smeets and González, 2000). The presence of dopamine in elasmobranch pallial neurons was reported in a skate with dopamine antibodies (Meredith and Smeets, 1987), which suggests that the pallial TH-ir cells of catshark are also dopaminergic, and thus they may co-release dopamine and SST. The presence of dopamine as end-product of TH in mammalian cortical interneurons is not known (see Asmus et al., 2007).

With regards the colocalization of TH immunoreactivity and somatostatin *PSST6* expression in other brain regions, we have not found similar populations in other brain regions excepting the lateral preoptic/suprachiasmatic area, despite the abundant co-distribution of TH-ir and *PSST1*+ or *PSST6*+ cells in regions such as the dorsal hypothalamus, posterior tubercle, ventral tegmental area and substantia nigra. In other vertebrates, a few examples of colocalization of TH and SST immunoreactivities have been reported in urodeles and rats. Low numbers of TH-ir neurons showed colocalization of catecholamines and SST in the ventral preoptic area of a urodele (González, Moreno, Morona, & López, 2003), but not in many brain regions where catecholaminergic neurons and SST-ir cells are largely co-distributed. In the rat, double-labeled TH-ir/SST-ir cells were frequent in the preoptic periventricular nucleus and double-labeled cells were also observed in the hypothalamus (Sakanaka, Magari, & Inoue, 1990). Thus, the presence of TH/SST double labeled neurons appears restricted to a few populations in the brain of vertebrates.

### Concluding remarks

The diversity of *PSST* genes reported in vertebrates (Tostivint et al., 2014; Tostivint, Gaillard, Mazan, & Pézeron, 2019), together with available comparative data of brain expression (see above) indicates that SSTs are involved in a number of neural circuits (Urban-Ciecko and Barth, 2016). The genes for six SST receptor types (SSTR1-6) have been cloned in vertebrates (Ocampo Daza, Sundström, Bergqvist, & Larhammar, 2012) and Northern blotting and *in situ* hybridization studies have shown that the transcripts of 5 receptors of rat are expressed in the CNS (Meyerhof, Wulfsen, Schönrock, Fehr, & Richter, 1992; Señarís, Humphrey, & Emson, 1994). SST exerts potent inhibitory actions on hormone secretion and neuronal excitability mediated by G protein-coupled receptors (Lin and Peter, 2001; Tostivint et al., 2014; Urban-Ciecko and Barth, 2016; Günther et al., 2018). Pharmacology studies reveal that properties of SSTRs differ with regards affinity to ligands SST-14 and SST-28, expression patterns and regulation (Lin and Peter, 2001; Günther et al., 2018). Moreover, phylogenetic analyses of the SSTR gene family reveal the early diversification of these genes in vertebrates, giving rise to five different SSTR (SSTRa-e) subtype genes in elasmobranchs (Ocampo Daza, Sundström, Bergqvist, & Larhammar, 2012). Taken together the diversity of SSTs and the wide regional differences in expression patterns of *PSSTs* in the catshark, it is expected that SSTRs will be also expressed differentially in the brain of elasmobranchs. Future studies characterizing the affinities of these receptors for the different somatostatin peptides generated in elasmobranchs, as well as characterization of brain distribution of five receptors may help to understand the variety of functions in which the somatostatinergic systems may be implicated.

## Acknowledgements

Grant sponsors: Spanish Ministry of Economy and Competitiveness and the European Regional Development Fund 2007-2013 (Grant number: BFU-2017-87079-P to MCR). Agence Nationale de la Recherche (ANR) grant NEMO no ANR-14-CE02-0020-01 (to HT).

## Abbreviations used in table 2 and figures

APV: Ventrolateral periventricular area
AU: Cerebellar auricle
BSA: Basal superficial area
CB: Cerebellum
Cc: Cerebellar crest
CCa: Central canal
cDP: Caudal region of dorsal pallium
CG: Central gray
Ch: optic chiasm
Cho: Choroid plexus
CN: Cerebellar nucleus
CSA: Central subpallial area
DH: Dorsal horn
DON: Dorsal octavolateralis nucleus
DP: Dorsal pallium
En: Entopeduncular nucleus
EW: Edinger-Westphal nucleus
gl: glomeruli
gr: Cerebellar granular layer
H: Hypophysis
Ha: Habenula
Hy: Hypothalamus
ica: Interstitial nucleus of the anterior commissure
IHL: Inferior hypothalamic lobe
III: Oculomotor nucleus
IP: Interpeduncular nucleus
IRF: Inferior reticular formation Ist Isthmus
IV: Trochlear nucleus
LC: Locus coeruleus
LMN: Lateral mesencephalic nucleus
LR: Lateral recess of pallium
LX: Vagal lobe
LP: Lateral pallium
mDP: Medial part of the dorsal pallium
ME: Median eminence
mi: mitral cell region
ml: Cerebellar molecular layer mlf Medial longitudinal fascicle
MON: Medial octavolateralis nucleus
MP: Medial pallium
MRF: Middle reticular formations
NLT: Lateral tuberal nucleus
NM: Median hypothalamic nucleus’
Nmlf: Nucleus of the medial longitudinal fascicle
Nr: Neuroporic recess
Nsm: Nucleus of the stria medullaris
NSV: Nucleus of the saccus vasculosus
OB: Olfactory bulb
OLA: Octavolateralis area
ON: Optic nerve
OT: Optic tectum P Pallium
pc: Posterior commissure
PO: Preoptic area
Pr: Posterior recess
PRO: Posterior recess organ
Pt: Pretectum
PTh: Prethalamus
PTN: Posterior tubercle nucleus
PTu: Posterior tubercle
PVO: Paraventricular organ
Rh: Rhombencephalon
SC: Spinal cord
SCh: Suprachiasmatic nucleus
Sco: Subcommissural organ
sDP: Superficial region of dorsal pallium
Se: Septal region
Sec: Caudal septum
Sep: Posterior medial septum
SN: Substantia nigra
Sp: Subpallium
SRF: Superior reticular formation
SRFl: Superior reticular formation, lateral part
SRFm: Superior reticular formation, medial part St Striatum
SV: Saccus vasculosus
Tg: Tegmentum
Th: Thalamus
trP: Tractus pallii
Vd: Trigeminal descending root and nucleus
VH: Ventral horn
VTA: Ventral tegmental area

## References

Adler, A., Zhao, R., Shin, M. E., Yasuda, R., & Gan, W. B. (2019). Somatostatin-expressing interneurons enable and maintain learning-dependent sequential activation of pyramidal neurons. Neuron, 102, 202–216.e7. doi: 10.1016/j.neuron.2019.01.036.

Adrio, F., Anadón, R., & Rodríguez-Moldes, I. (2008). Distribution of somatostatin immunoreactive neurons and fibres in the central nervous system of a chondrostean, the Siberian sturgeon (*Acipenser baeri*). Brain Research, 1209, 92–104. doi: 10.1016/j.brainres.2008.03.002.

Alvarez-Otero, R., Pérez, S.E., Rodríguez, M. A., Adrio, F., & Anadón, R. (1995). GABAergic neuronal circuits in the cerebellum of the dogfish *Scyliorhinus canicula* (Elasmobranchs): an immunocytochemical study. Neuroscience Letters, 187, 87–90. doi: 10.1016/0304-3940(95)11346-0

Alvarez-Otero, R., Perez, S. E., Rodriguez, M. A., & Anadón, R. (1996). Organisation of the cerebellar nucleus of the dogfish, *Scyliorhinus canicula* L.: a light microscopic, immunocytochemical, and ultrastructural study. Journal of Comparative Neurology, 368, 487–502. doi.org/10.1002/(SICI)1096-9861(19960513)368:4<487::AID-CNE2>3.0.CO;2-0.

Anadón, R., Molist. P., Rodríguez-Moldes, I., López, J. M., Quintela, I., Cerviño, M. C., Barja, P., & González, A. (2000). Distribution of choline acetyltransferase immunoreactivity in the brain of an elasmobranch, the lesser spotted dogfish (*Scyliorhinus canicula*). Journal of Comparative Neurology, 420, 139–170. doi: 10.1002/(SICI)1096-9861(19960513)368:4<487::AID-CNE2>3.0.CO;2-0

Anadón, R., Ferreiro-Galve, S., Sueiro, C., Graña, P., Carrera, I., Yáñez, J., & Rodríguez-Moldes, I. (2009). Calretinin-immunoreactive systems in the cerebellum and cerebellum-related lateral-line medullary nuclei of an elasmobranch, *Scyliorhinus canicula*. Journal of Chemical Neuroanatomy, 37, 46–54. doi: 10.1016/j.jchemneu.2008.09.003

Anadón, R., Molist, P., Pombal, M. A., Rodríguez-Moldes, I., & Rodicio, M. C. (1995). Marginal cells in the spinal cord of four elasmobranchs (*Torpedo marmorata*, *T. torpedo*, Raja undulata and Scyliorhinus canicula): evidence for homology with lamprey intraspinal stretch receptor neurons. European Journal of Neuroscience, 7, 934–43. doi.org/10.1111/j.1460-9568.1995.tb01081.x

Anadón, R., Rodríguez-Moldes, I., & Adrio, F. (2013). Glycine-immunoreactive neurons in the brain of a shark (*Scyliorhinus canicula* L.). Journal of Comparative Neurology, 521, 3057–3082. doi: 10.1002/cne.23332.

Anderson, S. A., Marín, O., Horn, C., Jennings, K., & Rubenstein, J. L. (2001). Distinct cortical migrations from the medial and lateral ganglionic eminences. Development, 128, 353–363. Retrieved from https://dev.biologists.org/content/128/3/353

Anglade, I., Zandbergen, T., & Kah, O. (1993). Origin of the pituitary innervation in the goldfish. Cell and Tissue Research, 273, 345–55. doi: 10.1007/BF00312837.

Armstrong, C. L., Chung, S. H., Armstrong, J. N., Hochgeschwender, U., Jeong, Y. G., & Hawkes, R. (2009). A novel somatostatin-immunoreactive mossy fiber pathway associated with HSP25-immunoreactive Purkinje cell stripes in the mouse cerebellum. Journal of Comparative Neurology, 517, 524–538. doi: 10.1002/cne.22167.

Asmus, S. E., Anderson, E. K., Ball, M. W., Barnes, B. A., Bohnen, A. M., Brown, A. M., Hartley, L. J., Lally, M. C., Lundblad, T. M., Martin, J. B., Moss, B. D., Phelps, K. D., Phillips, L. R., Quilligan, C. G., Steed, R. B., Terrell, S. L., & Warner, A. E. (2008). Neurochemical characterization of tyrosine hydroxylase-immunoreactive interneurons in the developing rat cerebral cortex. Brain Research, 1222, 95–105. doi: 10.1016/j.brainres.2008.05.053.

Barreiro-Iglesias, A., Mysiak, K. S., Adrio, F., Rodicio, M. C., Becker, C. G., Becker, T., & Anadón, R. (2013). Distribution of glycinergic neurons in the brain of glycine transporter-2 transgenic Tg(glyt2:Gfp) adult zebrafish: relationship to brain-spinal descending systems. Journal of Comparative Neurology, 521, 389–425. doi: 10.1002/cne.23179.

Becerra, M., Manso, M. J., Rodríguez-Moldes, I., & Anadón, R. (1995). Ontogeny of somatostatin-immunoreactive systems in the brain of the brown trout (Teleostei). Anatomy and Embryology (Berlin), 191, 119–137. doi.org/10.1007/BF00186784

Böhm, U. L., Prendergast, A., Djenoune, L., Nunes Figueiredo, S., Gomez, J., Stokes, C., Kaiser, S., Suster, M., Kawakami, K., Charpentier, M., Concordet, J. P., Rio, J. P., Del Bene, F., & Wyart, C. (2016). CSF-contacting neurons regulate locomotion by relaying mechanical stimuli to spinal circuits. Nature Communications, 7, 10866. doi: 10.1038/ncomms10866.

Boord, R. L., & Northcutt, R. G. (1988). Medullary and mesencephalic pathways and connections of lateral line neurons of the spiny dogfish *Squalus acanthias*. Brain Behavior and Evolution, 32, 76–88. doi: 10.1159/000116535.

Brazeau, P., Rivier, J., Vale, W., & Guillemin, R. (1974). Inhibition of growth hormone secretion in the rat by synthetic somatostatin. Endocrinology, 94, 184–187. doi:10.1210/endo-94-1-184

Buchanan, J. T., Brodin, L., Hökfelt, T., Van Dongen, P. A., & Grillner, S. (1987). Survey of neuropeptide-like immunoreactivity in the lamprey spinal cord. Brain Research, 408, 299–302. doi: 10.1016/0006-8993(87)90392-1

Cameron, A. A., Plenderleith, M. B., & Snow, P. J. (1990). Organization of the spinal cord in four species of elasmobranch fish: cytoarchitecture and distribution of serotonin and selected neuropeptides. Journal of Comparative Neurology, 297, 201–218. doi: 10.1002/cne.902970204

Canosa, L. F., Cerdá-Reverter, J. M., & Peter, R. E. (2004). Brain mapping of three somatostatin encoding genes in the goldfish. Journal of Comparative Neurology, 474, 43–57. doi: 10.1002/cne.20097.

Carrera, I., Sueiro, C., Molist, P., Ferreiro, S., Adrio, F., Rodríguez, M. A., Anadón, R., & Rodríguez-Moldes, I. (2005). Temporal and spatial organization of tyrosine hydroxylase-immunoreactive cell groups in the embryonic brain of an elasmobranch, the lesser-spotted dogfish *Scyliorhinus canicula*. Brain Research Bulletin, 66, 541–545. doi: 10.1016/j.brainresbull.2005.02.010

Carrera, I., Molist, P., Anadón, R., & Rodríguez-Moldes, I. (2008a). Development of the serotoninergic system in the central nervous system of a shark, the lesser spotted dogfish *Scyliorhinus canicula*. Journal of Comparative Neurology, 511, 804–831. doi: 10.1002/cne.21857.

Carrera, I., Ferreiro-Galve, S., Sueiro, C., Anadón, R., & Rodríguez-Moldes, I. (2008b). Tangentially migrating GABAergic cells of subpallial origin invade massivelythe pallium in developing sharks. Brain Research Bulletin, 75, 405–409. doi: 10.1016/j.brainresbull.2007.10.013.

Carrera, I., Anadón, R., & Rodríguez-Moldes, I. (2012). Development of tyrosine hydroxylase-immunoreactive cell populations and fiber pathways in the brain of the dogfish *Scyliorhinus canicula*: new perspectives on the evolution of the vertebrate catecholaminergic system. Journal of Comparative Neurology, 520, 3574–3603. doi: 10.1002/cne.23114.

Chiba, A., Honma, Y., Ito, S., & Homma, S. (1989). Somatostatin-immunoreactivity in the brain of the gommy shark, *Mustelus manazo* Bleeker, with special regard to the hypothalamo-hypophyseal system. Biomedical Research, 10, Supplement 3, 1–12.

Christenson, J., Alford, S., Grillner, S., & Hökfelt, T. (1991). Co-localized GABA and somatostatin use different ionic mechanisms to hyperpolarize target neurons in the lamprey spinal cord. Neuroscience Letters, 134, 93–97. doi: 10.1016/0304-3940(91)90516-v.

Coolen, M., Menuet, A., Chassoux, D., Compagnucci, C., Henry, S., Lévèque, L., Da Silva, C., Gavory, F., Samain, S., Wincker, P., Thermes, C., D’Aubenton-Carafa, Y., Rodríguez-Moldes, I., Naylor, G., Depew, M., Sourdaine, P., & Mazan, S. (2009). The dogfish *Scyliorhinus canicula*, a reference in jawed vertebrates. In R. R. Behringer, A. D. Johnson, R. E. Krumlauf (Eds.), Emerging Model Organisms. A Laboratory Manual (pp. 431–446). Vol. 1.Cold Spring Harbor, NY: Cold Spring Harbor Laboratory Press.

Cornide-Petronio, M. E., Anadón, R., Barreiro-Iglesias, A., & Rodicio, M. C. (2013). Serotonin 1A receptor (5-HT1A) of the sea lamprey: cDNA cloning and expression in the central nervous system. Brain Structure and Function, 218, 1317–1335. doi: 10.1007/s00429-012-0461-y.

El-Salhy, M. (1984). Immunocytochemical investigation of the gastro-entero-pancreatic (GEP) neurohormonal peptides in the pancreas and gastrointestinal tract of the dogfish *Squalus acanthias*. Histochemistry, 80, 193–205. MEDLINE PMID: 6370932.

Esclapez, M., & Houser, C. R. (1995). Somatostatin neurons are a subpopulation of GABA neurons in the rat dentate gyrus: evidence from colocalization of pre-prosomatostatin and glutamate decarboxylase messenger RNAs. Neuroscience, 64, 339–355. doi.org/10.1016/0306-4522(94)00406-U.

Ferreiro-Galve, S., Carrera, I., Candal, E., Villar-Cheda, B., Anadón, R., Mazan, S., & Rodríguez-Moldes, I. (2008). The segmental organization of the developing shark brain based on neurochemical markers, with special attention to the prosencephalon. Brain Research Bulletin, 75, 236–240. doi: 10.1016/j.brainresbull.2007.10.048

Ferreiro-Galve, S., Rodríguez-Moldes, I., & Candal, E. (2012). Pax6 expression during retinogenesis in sharks: comparison with markers of cell proliferation and neuronal differentiation. Journal of Experimental Zoology Part B: Molecular and Developmental Evolution, 318, 91–108. doi: 10.1002/jezb.21448.

Fiebig, E. (1988). Connections of the corpus cerebelli in the thornback guitarfish, *Platyrhinoidis triseriata* (Elasmobranchii): a study with WGA-HRP and extracellular granule cell recording. Journal of Comparative Neurology, 268, 567–583. doi: 10.1002/cne.902680407.

Filippi, A, Mahler, J, Schweitzer, J, & Driever, W. (2010). Expression of the paralogous tyrosine hydroxylase encoding genes th1 and th2 reveals the full complement of dopaminergic and noradrenergic neurons in zebrafish larval and juvenile brain. Journal of Comparative Neurology, 518, 423–438. doi: 10.1002/cne.22213.

Finger, T. E. (2008). Sorting food from stones: the vagal taste system in goldfish, *Carassius auratus*. Journal of Comparative Physiology A, 194, 135–143. doi: 10.1007/s00359-007-0276-0.

Fitzpatrick-McElligott, S., Card, J.P., Lewis, M. E., & Baldino, F. Jr. (1988). Neuronal localization of prosomatostatin mRNA in the rat brain with in situ hybridization histochemistry. Journal of Comparative Neurology, 273, 558–572. doi: 10.1002/cne.902730410.

Folgueira, M., Anadón, R., & Yáñez, J. (2006). Afferent and efferent connections of the cerebellum of a salmonid, the rainbow trout (*Oncorhynchus mykiss*): a tract-tracing study. Journal of Comparative Neurology, 497, 542–565. doi: 10.1002/cne.20979.

Gilland, E., Straka, H., Wong, T. W., Baker, R., & Zottoli, S. J. (2014). A hindbrain segmental scaffold specifying neuronal location in the adult goldfish, *Carassius auratus*. Journal of Comparative Neurology, 522, 2446–2464. doi: 10.1002/cne.23544.

Giuliani, A., Minelli, D., Quaglia, A., & Villani, L. (2002). Telencephalo-habenulo-interpeduncular connections in the brain of the shark *Chiloscyllium arabicum*. Brain Research, 926, 186–190. doi: 10.1016/s0006-8993(01)03310-8.

González, A., Moreno, N., Morona, R., & López, J. M. (2003). Somatostatin-like immunoreactivity in the brain of the urodele amphibian *Pleurodeles waltl*. Colocalization with catecholamines and nitric oxide. Brain Research, 965, 246–258. doi: 10.1016/s0006-8993(02)04210-5.

Grillner, S., Williams, T., & Lagerbäck, P. A. (1984). The edge cell, a possible intraspinal mechanoreceptor. Science, 223(4635), 500–503. doi: 10.1016/s0006-8993(02)04210-5.

Günther, T., Tulipano, G., Dournaud, P., Bousquet, C., Csaba, Z., Kreienkamp, H. J., Lupp, A., Korbonits, M., Castaño, J. P., Wester, H. J., Culler, M., Melmed, S., & Schulz, S. (2018). International Union of Basic and Clinical Pharmacology. CV. Somatostatin receptors: structure, function, ligands, and new nomenclature. Pharmacological Reviews, 70, 763–835. doi: 10.1124/pr.117.015388.

Harvey, A. R., Heavens, R. P., Yellachich, L. A., & Sirinathsinghji, D. J. (2001). Expression of messenger RNAs for glutamic acid decarboxylase, preprotachykinin, cholecystokinin, somatostatin, proenkephalin and neuropeptide Y in the adult rat superior colliculus. Neuroscience, 103, 443–455. doi.org/10.1016/S0306-4522(00)00581-9.

Higashijima, S., Mandel, G., & Fetcho, J. R. (2004). Distribution of prospective glutamatergic, glycinergic, and GABAergic neurons in embryonic and larval zebrafish. Journal of Comparative Neurology, 480, 1–18. doi: 10.1002/cne.20278.

Holmgren, S., & Nilsson, S. (1983). Bombesin-, gastrin/CCK-, 5-hydroxytryptamine-, neurotensin-, somatostatin-, and VIP-like immunoreactivity and catecholamine fluorescence in the gut of the elasmobranch, *Squalus acanthias*. Cell and Tissue Research, 234, 595–618. Retrieved from PubMed PMID: 6362887.

Holmqvist, B. I., & Ekström, P. (1995). Hypophysiotrophic systems in the brain of the Atlantic salmon. Neuronal innervation of the pituitary and the origin of pituitary dopamine and nonapeptides identified by means of combined carbocyanine tract tracing and immunocytochemistry. Journal of Chemical Neuroanatomy, 8, 125–145. doi.org/10.1016/0891-0618(94)00041-Q.

Hsu, Y. W., Wang, S. D., Wang, S., Morton, G., Zariwala, H. A., de la Iglesia, H. O., & Turner, E. E. (2014). Role of the dorsal medial habenula in the regulation of voluntary activity, motor function, hedonic state, and primary reinforcement. Journal of Neuroscience, 34, 11366–11384. doi: 10.1523/JNEUROSCI.1861-14.2014.

Huesa, G., Anadón, R., & Yáñez, J. (2003). Afferent and efferent connections of the cerebellum of the chondrostean *Acipenser baeri*: a carbocyanine dye (DiI) tracing study. Journal of Comparative Neurology, 460, 327–344. doi: 10.1002/cne.10629.

Inagaki, S., Shiosaka, S., Sekitani, M., Noguchi, K., Shimada, S., & Takagi, H. (1989). In situ hybridization analysis of the somatostatin-containing neuron system in developing cerebellum of rats. Brain Research Molecular Brain Research, 6, 289–295. doi.org/10.1016/0169-328X(89)90074-0.

Jalalvand, E., Robertson, B., Wallén, P., Hill, R. H., & Grillner, S. (2014). Laterally projecting cerebrospinal fluid-contacting cells in the lamprey spinal cord are of two distinct types. Journal of Comparative Neurology, 522, 1753–1768. doi: 10.1002/cne.23542.

Jalalvand, E., Robertson, B., Tostivint, H., Löw, P., Wallén, P., & Grillner, S. (2018). Cerebrospinal fluid-contacting neurons sense pH changes and motion in the hypothalamus. Journal of Neuroscience, 38, 7713–7724. doi: 10.1523/JNEUROSCI.3359-17.2018.

Johansson, O., Hökfelt, T., & Elde, R. P. (1984). Immunohistochemical distribution of somatostatin-like immunoreactivity in the central nervous system of the adult rat. Neuroscience, 13, 265–339. doi.org/10.1016/0306-4522(84)90233-1.

Johnston, S. A., & Maler, L. (1992). Anatomical organization of the hypophysiotrophic systems in the electric fish, *Apteronotus leptorhynchus*. Journal of Comparative Neurology, 317, 421–437. doi: 10.1002/cne.903170408.

Kinkhabwala, A., Riley, M., Koyama, M., Monen, J., Satou, C., Kimura, Y., Higashijima, S., & Fetcho, J. (2011). A structural and functional ground plan for neurons in the hindbrain of zebrafish. Proceedings of the National Academy of Sciences U S A, 108, 1164–1169. doi: 10.1073/pnas.1012185108.

Kiyama, H., & Emson, P. C. (1990). Distribution of somatostatin mRNA in the rat nervous system as visualized by a novel non-radioactive in situ hybridization histochemistry procedure. Neuroscience, 38, 223–244. doi.org/10.1016/0306-4522(90)90388-K.

Knowles, F. (1965). Evidence by a dual control, by neurosecretion, of hormone synthesis and hormone release in the pituitary of the dogfish, Scylliorhinus stellaris. Philosophical Transactions of the Royal Society B, 249, 435–456. doi.org/10.1098/rstb.1965.0018.

Lee, R. K., Eaton, R. C., & Zottoli, S. J. (1993). Segmental arrangement of reticulospinal neurons in the goldfish hindbrain. Journal of Comparative Neurology, 329, 539–556. https://doi.org/10.1002/cne.903290409

Lin, X., & Peter, R. E. (2001). Somatostatins and their receptors in fish. Comparative Biochemistry and Physiology. Part B, Biochemistry & Molecular Biology, 129, 543–550. https://doi.org/10.1016/S1096-4959(01)00362-1

Manso, M. J., & Anadon, R. (1991a). The optic tectum of the dogfish *Scyliorhinus canicula* L.: a Golgi study. Journal of Comparative Neurology, 307, 335–349. doi: 10.1002/cne.903070212.

Manso, M. J., & Anadón, R. (1991b). Specialized presynaptic dendrites in the stratum cellulare externum of the optic tectum of an elasmobranch, *Scyliorhinus canicula* L. Neuroscience Letters, 129, 291–293. doi: 10.1016/0304-3940(91)90483-a

Manso, M. J., & Anadón, R. (1993). Golgi study of the telencephalon of the small-spotted dogfish *Scyliorhinus canicula* L. Journal of Comparative Neurology, 333, 485–502. doi: 10.1002/cne.903330403.

Mellinger, J. C. A. (1964). Les relations neuro-vasculo-glandulaires dans l’appareil hypophysaire de la roussette, *Scyliorhinus caniculus* (L.) (Poissons Elasmobranches). These Sciences, Strasburg; Impr. Alsatia Colmar, 1963, 211 pp. Reimpr. in: Archives de Anatomie, Histologie et Embryologie, 47, 1–201.

Meredith, G. E., & Smeets, W. J. (1987). Immunocytochemical analysis of the dopamine system in the forebrain and midbrain of *Raja radiata*: evidence for a substantia nigra and ventral tegmental area in cartilaginous fish. Journal of Comparative Neurology, 265, 530–548. doi: 10.1002/cne.902650407.

Metcalfe, W. K., Mendelson, B., & Kimmel, C. B. (1986). Segmental homologies among reticulospinal neurons in the hindbrain of the zebrafish larva. Journal of Comparative Neurology, 251, 147–159. doi.org/10.1002/cne.902510202

Meurling, P., & Rodríguez, E. M. (1990). The paraventricular and posterior recess organs of elasmobranchs: A system of cerebrospinal fluid-contacting neurons containing immunoreactive serotonin and somatostatin. Cell and Tissue Research, 259, 463–473. doi.org/10.1007/BF01740772

Meurling, P., Rodríguez, E. M., Peña, P., Grondona, J. M., & Pérez, J. (1996). Hypophysial and extrahypophysial projections of the neurosecretory system of cartilaginous fishes: an immunocytochemical study using a polyclonal antibody against dogfish neurophysin. Journal of Comparative Neurology, 373, 400–421. doi: 10.1002/(SICI)1096-9861(19960923)373:3<400::AID-CNE6>3.0.CO;2-6.

Meyerhof, W., Wulfsen, I., Schönrock, C., Fehr, S., & Richter, D. (1992). Molecular cloning of a somatostatin-28 receptor and comparison of its expression pattern with that of a somatostatin-14 receptor in rat brain. Proceedings of the National Academy of Sciences U S A, 89, 10267–10271. doi: 10.1073/pnas.89.21.10267.

Molist, P., Rodriguez-Moldes, I., & Anadon, R. (1992). Immunocytochemical and electron-microscopic study of the elasmobranch nucleus sacci vasculosi. Cell and Tissue Research, 270, 395–404. doi.org/10.1007/BF00328023

Molist, P., Rodríguez-Moldes, I., & Anadón, R. (1993). Organization of catecholaminergic systems in the hypothalamus of two elasmobranch species, Raja undulata and Scyliorhinus canicula. A histofluorescence and immunohistochemical study. Brain Behavior and Evolution, 41, 290–302. doi: 10.1159/000113850.

Morita, Y., & Finger, T. E. (1985). Topographic and laminar organization of the vagal gustatory system in the goldfish, *Carassius auratus*. Journal of Comparative Neurology, 238, 187–201. doi: 10.1002/cne.902380206.

Northcutt, R. G., Reiner, A., & Karten, H. J. (1988). Immunohistochemical study of the telencephalon of the spiny dogfish, *Squalus acanthias*. Journal of Comparative Neurology, 277, 250–267. doi: 10.1002/cne.902770207.

Nozaki, M., Tsukahara, T., & Kobayashi, H. (1984). An immunocytochemical study on the distribution of neuropeptides in the brain of certain species of fish. Biomedical Research, 4, Supplementum, 135–145.

Ocampo Daza, D., Sundström, G., Bergqvist, C. A., & Larhammar, D. (2012). The evolution of vertebrate somatostatin receptors and their gene regions involves extensive chromosomal rearrangements. BMC Evolutionary Biology, 12, 231. doi: 10.1186/1471-2148-12-231.

Pose-Méndez, S., Candal, E., Adrio, F., & Rodríguez-Moldes, I. (2014). Development of the cerebellar afferent system in the shark *Scyliorhinus canicula*: insights into the basal organization of precerebellar nuclei in gnathostomes. Journal of Comparative Neurology, 522, 131–68. doi: 10.1002/cne.23393.

Priestley, J.V., Réthelyi, M., & Lund, P.K. (1991). Semi-quantitative analysis of somatostatin mRNA distribution in the rat central nervous system using in situ hybridization. Journal of Chemical Neuroanatomy, 4, 131–153. doi.org/10.1016/0891-0618(91)90037-D.

Puelles, L., & Rubenstein, J. L. (2003). Forebrain gene expression domains and the evolving prosomeric model. Trends in Neuroscience, 26, 469–476. doi: 10.1016/S0166-2236(03)00234-0.

Quan, F. B., Kenigfest, N. B., Mazan, S., & Tostivint, H. (2013). Molecular cloning of the cDNAs encoding three somatostatin variants in the dogfish (*Scylorhinus canicula*). General and Comparative Endocrinology, 180, 1–6. doi: 10.1016/j.ygcen.2012.10.007.

Quina, L.A., Harris, J., Zeng, H., & Turner, E. E. (2017). Specific connections of the interpeduncular subnuclei reveal distinct components of the habenulopeduncular pathway. Journal of Comparative Neurology, 525, 2632–2656. doi: 10.1002/cne.24221.

Quintana-Urzainqui, I., Sueiro, C., Carrera, I., Ferreiro-Galve, S., Santos-Durán, G., Pose-Méndez, S., Mazan, S., Candal, E., & Rodríguez-Moldes, I. (2012). Contributions of developmental studies in the dogfish *Scyliorhinus canicula* to the brain anatomy of elasmobranchs: insights on the basal ganglia. Brain Behavior and Evolution, 80, 127–141. doi: 10.1159/000339871.

Quintana-Urzainqui, I., Rodríguez-Moldes, I., Mazan, S., & Candal, E. (2015). Tangential migratory pathways of subpallial origin in the embryonic telencephalon of sharks: evolutionary implications. Brain Structure and Function, 220, 2905–2926. doi: 10.1007/s00429-014-0834-5.

Repérant, J., Miceli, D., Rio, J. P., Peyrichoux, J., Pierre, J., & Kirpitchnikova, E. (1986). The anatomical organization of retinal projections in the shark *Scyliorhinus canicula* with special reference to the evolution of the selachian primary visual system. Brain Research, 396, 227–248. doi: 10.1016/0165-0173(86)90013-5.

Rodríguez-Moldes, M. I. (1986). Estudio Ultraestructural e Histofluorescente del Hipotálamo de la Pintarroja (Scyliorhinus canicula L.) con Especial Referencia a los Sistemas Licor-Contactantes y al Saco Vasculoso. Doctoral Thesis. Universidad de Santiago de Compostela.

Rodríguez-Moldes, I. (2009). A developmental approach to forebrain organization in elasmobranchs: new perspectives on the regionalization of the telencephalon. Brain Behavior and Evolution, 74, 20–29. doi: 10.1159/000229010

Rodríguez-Moldes, I., & Anadón, R. (1987). Aminergic neurons in the hypothalamus of the dogfish, Scyliorhinus canicula L. (Elasmobranch). A histofluorescence study. Journal für Hirnforschung, 28, 685–693.

Rodríguez-Moldes, I., Scheuermann, D. W., Adriaensen, D., De Groodt-Lasseel, M. H., Molist, P., & Anadón, R. (1993). Microspectrofluorimetric study of monoamines in the hypothalamus of *Scyliorhinus stellaris* L. Journal für Hirnforschung, 34, 57–61.

Rodríguez-Moldes, I., Carrera, I., Pose-Méndez, S., Quintana-Urzainqui, I., Candal, E., Anadón, R., Mazan, S., & Ferreiro-Galve, S. (2011). Regionalization of the shark hindbrain: a survey of an ancestral organization. Frontiers in Neuroanatomy, 5, 16. doi: 10.3389/fnana.2011.00016. eCollection 2011.

Rodríguez-Moldes, I., Santos-Durán, G. N., Pose-Méndez, S., Quintana-Urzainqui, I., & Candal, E. (2017). The brains of cartilaginous fishes. In J. Kaas (Ed.), Evolution of Nervous Systems, 2nd edition. vol. 1 (pp. 77–97). Oxford: Elsevier.

Sakanaka, M., Magari, S., & Inoue, N. (1990). Somatostatin co-localizes with tyrosine hydroxylase in the nerve cells of discrete hypothalamic regions in rats. Brain Research, 516, 313–317. doi: 10.1016/0006-8993(90)90933-3.

Señarís, R. M., Humphrey, P. P., & Emson, P. C. (1994). Distribution of somatostatin receptors 1, 2 and 3 mRNA in rat brain and pituitary. European Journal of Neuroscience, 6, 1883–1896. doi.org/10.1111/j.1460-9568.1994.tb00579.x.

Smeets, W. J. (1981). Retinofugal pathways in two chondrichthyans, the shark *Scyliorhinus canicula* and the ray *Raja clavata*. Journal of Comparative Neurology, 195, 1–11. doi: 10.1002/cne.901950103.

Smeets, W. J. A. J. (1982). The afferent connections of the tectum mesencephali in two chondrichthyans, the shark *Scyliorhinus canicula* and the ray *Raja clavata*. Journal of Comparative Neurology, 205, 139–152. doi: 10.1002/cne.902050205

Smeets, W. J., & González, A. (2000). Catecholamine systems in the brain of vertebrates: new perspectives through a comparative approach. Brain Research Brain Research Reviews, 33, 308–79. doi.org/10.1016/S0165-0173(00)00034-5.

Smeets, W. J. A. J., Nieuwenhuys, R., & Roberts, B. L. (1983). The Central Nervous System of Cartilaginous Fishes. Berlin, Heidelberg, New York: Springer.

Smeets, W. J., & Timerick, S. J. (1981). Cells of origin of pathways descending to the spinal cord in two chondrichthyans, the shark *Scyliorhinus canicula* and the ray *Raja clavata*. Journal of Comparative Neurology, 202, 473–491. doi: 10.1002/cne.902020403

Stoff, J. S., Rosa, R., Hallac, R., Silva, P., & Epstein, F. H. (1979). Hormonal regulation of active chloride transport in the dogfish rectal gland. American Journal of Physiology, 237, F138–44. doi: 10.1152/ajprenal.1979.237.2.F138

Stuesse, S. L., Cruce, W. L., & Northcutt, R. G. (1994). Localization of catecholamines in the brains of Chondrichthyes (cartilaginous fishes). In W.J.A.J. Smeets, & A. Reiner, (Eds.), Phylogeny and development of catecholamine systems in the CNS of vertebrates (pp 21–47). Cambridge, UK: Cambridge University Press.

Sueiro, C., Carrera, I., Rodríguez-Moldes, I., Molist, P., & Anadón, R. (2003). Development of catecholaminergic systems in the spinal cord of the dogfish *Scyliorhinus canicula* (Elasmobranchs). Brain Research Developmental Brain Research, 142, 141–150. doi.org/10.1016/S0165-3806(03)00062-2.

Sueiro, C., Carrera, I., Molist, P., Rodríguez-Moldes, I., & Anadón, R. (2004). Distribution and development of glutamic acid decarboxylase immunoreactivity in the spinal cord of the dogfish *Scyliorhinus canicula* (elasmobranchs). Journal of Comparative Neurology, 478, 189–206. doi: 10.1002/cne.20285.

Sueiro, C., Carrera, I., Ferreiro, S., Molist, P., Adrio, F., Anadón, R., & Rodríguez-Moldes, I. (2007). New insights on saccus vasculosus evolution: a developmental and immunohistochemical study in elasmobranchs. Brain Behavior and Evolution, 70, 187–204. https://doi.org/10.1159/000104309

Teijido, O., Manso, M. J., & Anadón, R. (2002). Distribution of thyrotropin-releasing hormone immunoreactivity in the brain of the dogfish *Scyliorhinus canicula*. Journal of Comparative Neurology, 454, 65–81. doi: 10.1002/cne.10431.

Timerick, S. J., Roberts, B. L., & Paul, D. H. (1992). Brainstem neurons projecting to different levels of the spinal cord of the dogfish *Scyliorhinus canicula*. Brain Behavior and Evolution, 39, 93–100. doi: 10.1159/000114107

Tostivint, H., Lihrmann, I., Bucharles, C., Vieau, D., Coulouarn, Y., Fournier, A., Conlon, J. M., & Vaudry, H. (1996). Occurrence of two somatostatin variants in the frog brain: characterization of the cDNAs, distribution of the mRNAs, and receptor-binding affinities of the peptides. Proceedings of the National Academy of Sciences U S A, 93, 12605–12610. doi: 10.1073/pnas.93.22.12605.

Tostivint, H., Dettaï, A., Quan, F. B., Ravi, V., Tay, B. H., Rodicio, M.C., Mazan, S., Venkatesh, B., & Kenigfest, N. B. (2016). Identification of three somatostatin genes in lampreys. General and Comparative Endocrinology, 237, 89–97. doi: 10.1016/j.ygcen.2016.08.006.

Tostivint, H., Ocampo Daza, D., Bergqvist, C. A., Quan, F. B., Bougerol, M., Lihrmann, I., & Larhammar, D. (2014). Molecular evolution of GPCRs: Somatostatin/urotensin II receptors. Journal of Endocrinology, 52, T61–86. doi: 10.1530/JME-13-0274.

Tostivint, H., Quan, F. B., Bougerol, M., Kenigfest, N. B., & Lihrmann, I. (2013). Impact of gene/genome duplications on the evolution of the urotensin II and somatostatin families. General and Comparative Endocrinology, 188, 110–117. doi: 10.1016/j.ygcen.2012.12.015.

Tostivint, H., Gaillard, A.L., Mazan, S., & Pézeron, G. (2019). Revisiting the evolution of the somatostatin family: Already five genes in the gnathostome ancestor. General and Comparative Endocrinology, 279, 139–147. doi: 10.1016/j.ygcen.2019.02.022.

Trabucchi, M., Tostivint, H., Lihrmann, I., Jégou, S., Vallarino, M., & Vaudry, H. (1999). Molecular cloning of the cDNAs and distribution of the mRNAs encoding two somatostatin precursors in the African lungfish *Protopterus annectens*. Journal of Comparative Neurology, 410, 643–652. doi.org/10.1002/(SICI)1096-9861(19990809)410:4<643::AID-CNE10>3.0.CO;2-%23.

Trabucchi, M., Tostivint, H., Lihrmann, I., Sollars, C., Vallarino, M., Dores, R. M., & Vaudry, H. (2002). Polygenic expression of somatostatin in the sturgeon *Acipenser transmontanus*: molecular cloning and distribution of the mRNAs encoding two somatostatin precursors. Journal of Comparative Neurology, 443, 332–345. doi.org/10.1002/cne.10126.

Trabucchi, M., Tostivint, H., Lihrmann, I., Blähser, S., Vallarino, M., & Vaudry, H. (2003). Characterization of the cDNA encoding a somatostatin variant in the chicken brain: comparison of the distribution of the two somatostatin precursor mRNAs. Journal of Comparative Neurology, 461, 441–451. doi: 10.1002/cne.10690.

Turner, K. J., Hawkins, T. A., Yáñez, J., Anadón, R., Wilson, S. W., & Folgueira, M. (2016). Afferent connectivity of the zebrafish habenulae. Frontiers in Neural Circuits, 10, 30. doi: 10.3389/fncir.2016.00030.

Urban-Ciecko, J., & Barth, A. L. (2016). Somatostatin-expressing neurons in cortical networks. Nature Reviews Neuroscience, 17, 401–409. doi: 10.1038/nrn.2016.53.

Villar-Cerviño, V., Holstein, G. R., Martinelli, G. P., Anadón, R., & Rodicio, M. C. (2008). Glycine-immunoreactive neurons in the developing spinal cord of the sea lamprey: comparison with the gamma-aminobutyric acidergic system. Journal of Comparative Neurology, 508, 112–130. doi: 10.1002/cne.21661.

Vincent, S. R., McIntosh, C. H., Buchan, A. M., & Brown, J. C. (1985). Central somatostatin systems revealed with monoclonal antibodies. Journal of Comparative Neurology, 238, 169–186. https://doi.org/10.1002/cne.902380205

Wang, M., & Bradley, R. M. (2010). Properties of GABAergic neurons in the rostral solitary tract nucleus in mice. Journal of Neurophysiology, 103, 3205–18. doi: 10.1152/jn.00971.2009.

Wilson, J. F., & Dodd, J. M. (1973). Distribution of monoamines in the diencephalon and pituitary of the dogfish, Scyliorhinus canicula L. Zeitschrift für Zellforschung und mikroskopische Anatomie, 137, 451–469. doi.org/10.1007/BF00307223

Wright, G. M. (1986). Immunocytochemical demonstration of growth hormone, prolactin and somatostatin-like immunoreactivities in the brain of larval, young adult and upstream migrant adult sea lamprey, *Petromyzon marinus*. Cell and Tissue Research, 246, 23–31. doi.org/10.1007/BF00218994

Yamamoto, K., Ruuskanen, J. O., Wullimann, M. F., & Vernier, P. (2010). Two tyrosine hydroxylase genes in vertebrates. New dopaminergic territories revealed in the zebrafish brain. Molecular and Cellular Neuroscience, 43, 394–402. doi: 10.1016/j.mcn.2010.01.006.

Yañez, J., & Anadón, R. (1996). Afferent and efferent connections of the habenula in the rainbow trout (*Oncorhynchus mykiss*): an indocarbocyanine dye (DiI) study. Journal of Comparative Neurology, 372, 529–543. doi: 10.1002/(SICI)1096-9861(19960902)372:4<529::AID-CNE3>3.0.CO;2-6.

Yáñez, J., Rodríguez, M., Pérez, S., Adrio, F., Rodríguez-Moldes, I., Manso, M. J., & Anadón, R. (1997). The neuronal system of the saccus vasculosus of trout (*Salmo trutta fario* and *Oncorhynchus mykiss*): an immunocytochemical and nerve tracing study. Cell and Tissue Research, 288, 497–507. doi.org/10.1007/s004410050835

Yáñez, J., Rodríguez-Moldes, I., & Anadón, R. (1992). Distribution of somatostatin-immunoreactivity in the brain of the larval lamprey (*Petromyzon marinus*). Journal of Chemical Neuroanatomy, 5, 511–520. doi.org/10.1016/0891-0618(92)90006-C.

Yavorska, I., & Wehr, M. (2016). Somatostatin-expressing inhibitory interneurons in cortical circuits. Frontiers in Neural Circuits, 10, 76. doi: 10.3389/fncir.2016.00076.

